# Flexible Cell Modeling Using Frequency Modulation

**DOI:** 10.1101/2024.05.03.592350

**Authors:** Jerry Jacob, Nitish Patel, Sucheta Sehgal

## Abstract

Computational models of the cell can be used to study the impact of drugs and assess pathological risks. Typically, computational models are computationally demanding or difficult to implement in dedicated hardware for real-time emulation. A new Frequency Modulation (FM) model is proposed to address these limitations. This new model utilizes a single sine generator with constant amplitude, but phase/frequency is modulated to emulate an action potential (AP). The crucial element of this model is the identification of the modulating signal. Focusing on FPGA implementation, we have utilized a piecewise linear polynomial with a fixed number of breakpoints to serve as a modulating signal. The ability to adapt this modulating signal permits the emulation of dynamic properties and coupling of cells. We have introduced a state controller that handles both of these requirements. The building blocks of the FM model have direct integer equivalents and are amenable to implementation on digital platforms like Field Programming Gate Arrays (FPGA).

We have demonstrated wavefront propagation of our model in 1-D and 2-D models of a tissue. Various parameters were used to quantify the wavefront propagation in 2-D tissues. We also emulate some cellular dysfunctions. The FM model can replicate any detailed cell model and emulate its corresponding tissue model. Overall, the results depict that the FM model has the potency for real-time cell and tissue emulation on an FPGA.

## Introduction

Mathematical models incorporating electrical functions have been pivotal in comprehending cellular mechanisms. The Hodgkin-Huxley (HH) model [15] was a pioneering example, accurately describing action potentials (APs) of a squid axon. Afterward, various other models like FitzHugh Nagumo (FHN) [10], Fenton Karma (FK) [9], Luo-Rudy-I (LR-I) [8], Beeler Reuter (BR) [27], Fabbri et al. [11] and O’Hara [29] models, have expanded on the HH model in many spheres. There exist other cardiac cell models based on purkinje fibres [30–33], ventricular nodes [8, 9, 27, 29] [34–51], sinoatrial node [11] [52, 53] and atrioventricular nodes [54]. These models have approximately 2-100 variables and many ordinary differential equations (ODEs). Therefore, implementing these models on an FPGA requires more computational space.

To overcome this issue, Sehgal et.al. [1] [2] formulated the Resonant Model (RM), which utilizes the Fourier Series (FS) to reconstruct various detailed models like Kurata et al. [14], Fabbri [11], Fenton Karma [9], O’Hara [29], Courtemanche et al. [12] and Dobrzynski et al. [13] models. The optimization of the Fourier Series coefficients was done by using the Levenberg-Marquardt algorithm [16]. The researchers exhibited the model’s ability to precisely emulate AP morphologies by altering the funny current (*I*_*f*_) block in the Fabbri et al. [11] model using linear and mixed fit-type equations. Jacob et al. [3] expanded upon this model by successfully applying the FS-based approach using non-linear regression [17] [18] to reconstruct APDs for the FHN, FK, BR, and LR-I models. They also quantified the resource usage of RM cells, demonstrating their lower resource requirements compared to Direct Digital Synthesis (DDS) methods. They [28] utilized the diffusion equation to simulate a ventricular tissue model using the RM.

Another emerging model that can be used to reconstruct the APDs is the Frequency Modulation Mobius (FMM) model [5] [6] [7]. Researchers have developed an R-software-based package for implementing the FMM [4], showcasing its effectiveness in reconstructing various data, including Iqgap gene expression, electrocardiography (ECG) records, and APs of the HH model.

The RM and FMM require multiple sinusoids to reconstruct major AP characteristics. The FM model attempts to reduce this aspect of the reconstruction. It simplifies the reconstruction of action potential durations (APDs) using a single sine wave. Hence, it is a mono-component model. It aims to utilize a sine wave with varying frequency but with a constant amplitude and phase. This study explores the efficacy of this model in terms of RMSE and *R*^2^ values for the reconstructions of the APDs of FitzHugh Nagumo (FHN) [10], Fenton Karma (FK), Beeler-Reuter and Fabbri [11] models. This paper also delves into the feasibility of parameterization, as in Sehgal et al. [1] for varying *I*_*f*_ block. We have designed a state controller for the FM model to emulate more dynamic properties to the replicated AP profiles. Utilizing this state controller, we created various propagation wavefronts for 1-D and 2-D tissues using O’Hara [29], FHN, and Fabbri [11] models. We could justify the wavefront propagation for a 2-D tissue using various metrics. Lastly, we introduced some cellular dysfunctions to the 2-D tissues and observed the alterations in the parameters.

### Theoretical Background

#### Concept

A Voltage Controlled Oscillator (VCO) is a device that outputs a repetitive signal with a frequency that depends on an input signal. It is typically used in broadcast communication, modems, PLLs and clock generation. Fig. 1 (A) shows, conceptually, how we intend to use our model. Fig. 1 (B) shows three signals: a pure sine wave as a reference, an example input signal and the resulting output of the VCO. The input has 4 segments: a negative step, a positive step, a slow negative ramp and a small negative constant. The frequency of *V*_*m*_ is smaller for the negative step and larger for the positive step. The frequency decreases slowly during the slow negative ramp; hence, the amplitude profile looks non-sinusoidal. As the input reduces and reaches a constant value, the frequency of the modulated sine wave remains constant. By astutely varying the profile like in Fig. 1(B), the APDs can be reconstructed.

**Fig 1.**
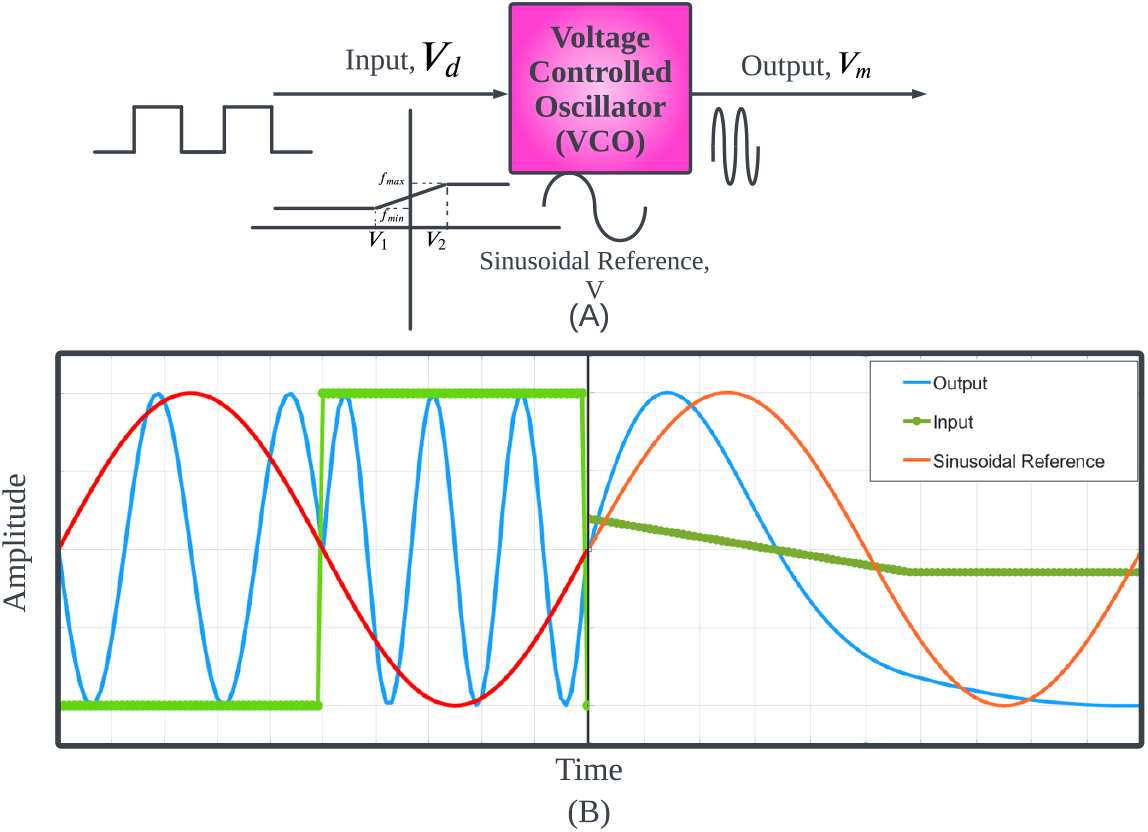
(A) Block diagram eliciting the working of a VCO (B) A sinusoidal wave of two periods altered by firstly a square wave and a combination of ramp and step functions.

#### Mathematical formulation of the FM model

A sinusoidal waveshape with a variation of phase, *ϕ*_*t*_(*t*) w.r.t a phase, *ϕ* (in radians) can be generated as in Eq 1.

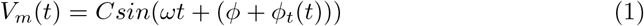

where C is the amplitude, *V*_*m*_(*t*) is the sinusoidal signal generated, *ω* is the constant frequency in radians/s, t is the time in milliseconds (s) where *t >* 0 and *ϕ*_*t*_(*t*) is the varying phase of the sinusoidal waveshape. *ϕ*_*t*_(*t*) can be obtained from Eq 2.

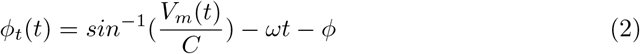

If Eq 1 is represented as the sinusoidal waveshape, which varies in terms of phase, *ϕ*_*t*_(*t*), then the frequency modulating factor, Δ_*w*_ (t) can vary the frequency as shown below in Eq 4. We can represent *ϕ*_*t*_(*t*) (in radians) in terms of varying frequency (in rad/s) as in Eq 3.

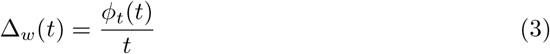

where *t >* 0 in s and Δ_*w*_ (t) is the frequency modulating factor.

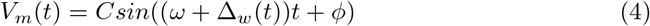

We can represent Eq.1 as in Eq 5.

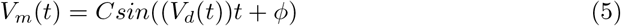

where *V*_*d*_(*t*) = (*ω* + Δ_*w*_(*t*)), t is time in s, *ϕ* is the constant phase (in radians), and *V*_*m*_(*t*) is the generated sinusoidal function in terms of varying frequency (in rad/s). Eq 5 depicts the VCO where *V*_*d*_(*t*) is introduced to get *V*_*m*_(*t*) as in Fig 1.

#### Test Case 1

This model relies on an accurate determination of Δ_*w*_. However, it is typically non-linear. Since Δ_*w*_(*t*) is updated at every time sample, this would imply that Δ_*w*_ will be a large 1-D matrix (12000 x 1) with a dimension proportional to the samples in one period. It is not feasible for a digital platform. A tractable solution is a set of linear piecewise equations fitted from the non-linear profile of Δ_*w*_.

This section demonstrates our identification methodology for our objective modulating signal. We intentionally choose three linear test signals (objective modulating signals) of Δ_*w*_(t) as shown in Fig. 2 (A),(B) and (C) with 12000 samples per period. Six linear equations with 7 breakpoints must be obtained for these test cases.

**Fig 2.**
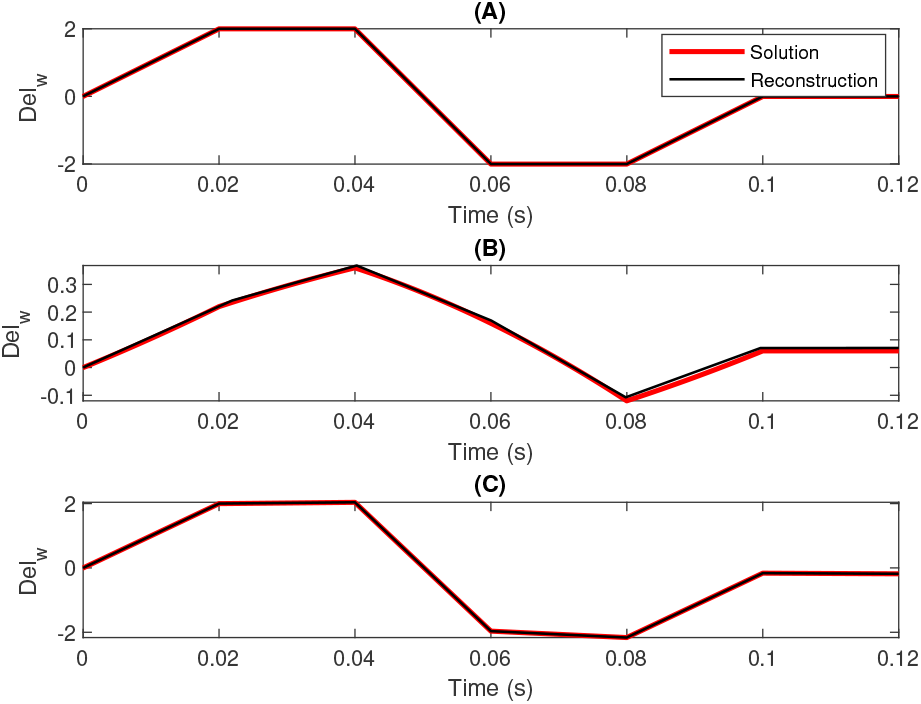
Utilization of three initial test signals (A) Signal 1, (B) Signal 2, and (C) Signal 3 used to evaluate the efficacy of the optimization technique of the novel FM model.

Piecewise linear equations offer the simplest implementation of FPGAs. The optimization of the coefficients of these equations is based on the sum of square error (SSE), the root mean square error (RMSE) and the coefficient of determination (*R*^2^). These are formulated in Eq 6, 7 and 8.

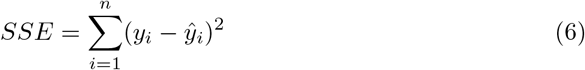

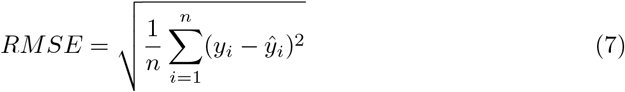

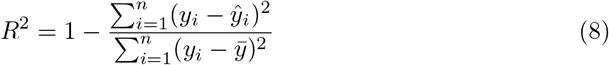

where *n* represents the total number of samples, *y*_*i*_ denotes the observed values, *ŷ*_*i*_ represents the predicted values, and 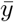 represents the mean of the observed values. The SSE values for all three signals (in Table.1) are below 0.9. Signals 1 and 3 portray a significantly reduced SSE of approximately 0.02. Regarding RMSE, Signal 2 is relatively high compared to its trapezoidal counterparts. The *R*^2^ values for all three signals exceed 0.99. These results demonstrate that the reconstructed piecewise linear polynomial generated using this technique can reconstruct the signal with greater precision.

**Table 1.** The sum of square errors (SSE), root mean square (RMSE) and *R*^2^ values of Signals 1,2 and 3.

| Signal | SSE | RMSE | $R^2$ value |
| --- | --- | --- | --- |
| Signal 1 | 0.0241 | 0.0014 | 1.0000 |
| Signal 2 | 0.9602 | 0.0089 | 0.9957 |
| Signal 3 | 0.0201 | 0.0012 | 1.0000 |

For realistic APDs, as discussed in later sections, Δ_*w*_ is non-linear and will require additional considerations. In addition, this method also facilitates the incorporation of the dynamic properties of APDs.

#### Test Case 2

Here, we consider non-linear modulating signals. We have specifically chosen sinusoids as they offer an additional metric total harmonic distortion (THD) along with RMSE to comprehend the efficiency of fitted waveshape to the sine waves. THD is formulated in Eq 9.

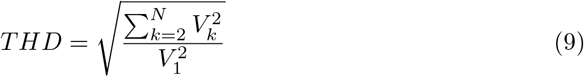

where *V*_*k*_ represents the harmonic components from k=2,., N and *V*_1_ represents the fundamental component of the signal.

Table.2 considers nine Δ_*w*_(t) profiles, sinusoids with varied frequencies and amplitudes. Twenty-three linear piecewise segments (24 breakpoints) were considered for all sine waves. For realistic APDs, we require 15-30 linear segments, so we chose 23 segments for this test case.

**Table 2.** The parameters of the tested sine waves in Test Case 2.

| Sine Signal | Frequency (rad/sec) | Amplitude (mV) | Total Harmonic Distortion (THD) | RMS error (RMSE) |
| --- | --- | --- | --- | --- |
| Sine Signal 1 | 2.0102 | 42.5175 | 0.0091 | 0.1286 |
| Sine Signal 2 | 2.2336 | 42.5175 | 0.0108 | 0.1059 |
| Sine Signal 3 | 4.0439 | 42.5271 | 0.0179 | 0.1425 |
| Sine Signal 4 | 5.1015 | 42.5271 | 0.0217 | 0.2496 |
| Sine Signal 5 | 6.6406 | 42.5315 | 0.0258 | 0.1923 |
| Sine Signal 6 | 7.7599 | 42.6564 | 0.0210 | 0.1833 |
| Sine Signal 7 | 8.5969 | 42.6978 | 0.0433 | 0.3630 |
| Sine Signal 8 | 9.2352 | 42.7750 | 0.0494 | 0.4390 |
| Sine Signal 9 | 10.1587 | 42.7970 | 0.0273 | 0.6461 |

The THD remains consistently within the range of approximately 0.01 to 0.05 despite variations in amplitudes and frequencies in the sine waves. The RMSE varies from 0.1 to 0.6 in all nine cases. These metrics affirm that the proposed technique can effectively reconstruct a sine wave signal.

#### Methodology

The methodology for the novel FM model is portrayed through a set of steps, as showcased in Fig. 3(A) and Fig. 3(B):

**Fig 3.**
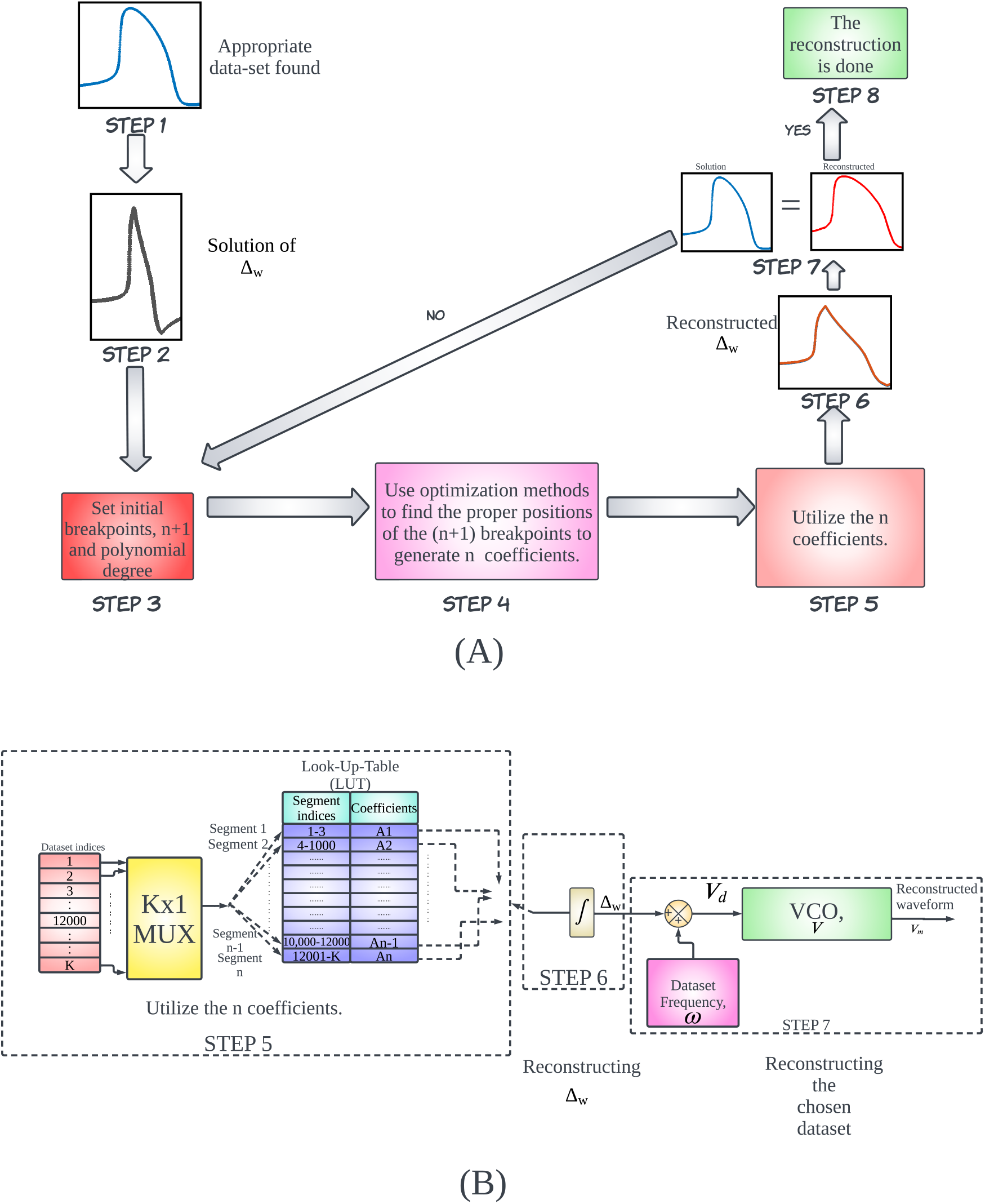
(A) Overall methodology of the novel FM model (B) Utilization of various polynomial segments using integrators to get the reconstructed waveform.

Step 1. Normalize the APD (time series of length K) to a custom numeric range.

Step 2. Generating a Δ_*w*_ waveshape from the acquired dataset.

Step 3. Initializing the number of breakpoints (n+1) for n segments with a chosen polynomial degree.

Step 4. Utilizing an optimization method [19] [20] [21] [22] for linear piecewise fitting, deploying the MATLAB function fmincon [23] [26], to identify the optimum position of the n+1 breakpoints and the coefficients of each line segment. fmincon uses an algorithm that minimizes the Sum of Square Errors (SSE).

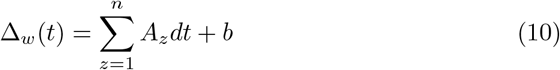

Step 5. Use the coefficients A1, A2, ……, An-1,An to create a Look-Up-Table (LUT).

Step 6. Using an integrator as per Eq 10, Δ_*w*_ is reconstructed by utilizing the segmented coefficients (as depicted in Fig. 3(B)). Δ_*w*_ is then added with the frequency of the data set, *ω*, to get *V*_*d*_, and fed into a VCO generating sinusoidal signal, V, to obtain the reconstructed waveform of the dataset, *V*_*m*_.

Step 7. Comparison of the reconstructed dataset to its original. If the comparison is not satisfactory, go back to step 3. If not, proceed to step 8.

Step 8. The reconstruction is successful.

#### Cell coupling using 1-D and 2-D diffusion equation

The simplest computational approach to create a 1-D cell coupling is known as centered second difference [56]. The equation can be represented as

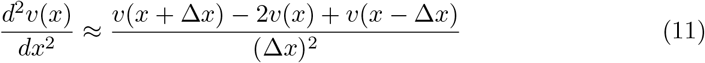

This formulation in Eq11 can be justified by combining the Taylor Series expansion for *v*(*x* + Δ*x*) and *v*(*x −* Δ*x*) (Strauss [55]). Consider the mesh size of the spacial variable as Δ*x*, then the *j*^*th*^ element of the array, v, *v*_*j*_ is of the value v for *x* = *j*Δ*x* represented by the equation below:

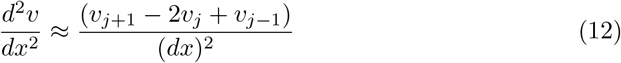

Using the expansion in Eq12, each cell can be coupled together along with a diffusion constant D to form D(*d*^2^*v*/*dx*^2^), which can be approximated as in Eq13.

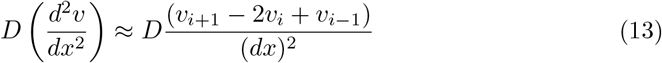

For simplicity of computation, Δ*x* is considered as 1. So Eq13 can be represented in Eq14.

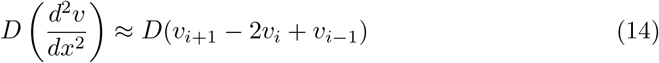

The following approach can be adapted to form a 2-D cell with a mesh size of Δ*x* and Δ*y* where Δ*x* is the mesh size in the x-axis and Δ*y* is the mesh size in the y-axis. For ease of computation, both the mesh sizes are considered the same, i.e.,Δ*x*=Δ*y*. The two-dimensional diffusion equation can be represented as

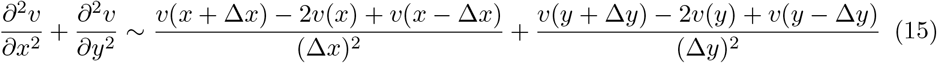

Consider the mesh sizes of the spacial variables to be Δ*x* and Δ*y*, then the *i*^*th*^ and *j*^*th*^ element of the array, *v*_*i*_ is of the value v for *x* = *i*Δ*x* and *v*_*j*_ is of the value v for *y* = *j*Δ*y*. Hence, we get Eq16,

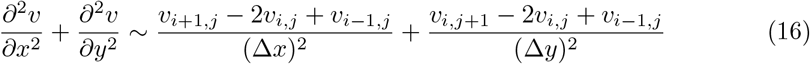

Using Eq16 along with the diffusion constant, D, we get Eq17.

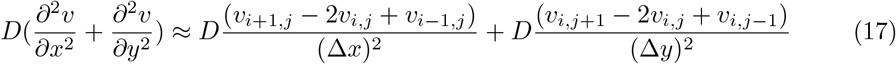

As mentioned earlier, for simplicity of calculation, Δ*x*=Δ*y*=1. By making this adjustment, we get Eq18,

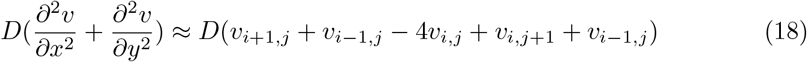

### Cell models used for the novel FM model

Post the successful reconstruction results obtained from various test signals, we proceed to apply the formulated approach for reconstructing several biophysical models, like the Beeler-Reuter (BR) [27], Fenton Karma (FK) [9], FitzHugh Nagumo (FHN) [10] and Fabbri et al. [11] models. The aim is to reconstruct the APDs of these models with high precision and fidelity using the proposed FM model.

The Beeler Reuter(BR) [27] model describes the ventricular cells in generic form. This model comprises 8 non-linear ordinary differential equations (ODEs). Six equations capture the gated channel’s state. The other two equations show the intracellular *Ca*^2+^ concentration and membrane voltage (*V*_*m*_). The slow inward current (*i*_*s*_) in the Beeler Reuter (BR) model emphasizes the plateau region formed by calcium ions in non-pacemaker cells. This model includes four distinct currents within its equations: 1) the initial fast current, *I*_*Na*_ 2) a slow inward current due to calcium ions; 3) a time-activated outward current, *I*_*x*1_; and 4) a time-dependent potassium current, *I*_*k*1_. The Fenton Karma (FK) [9] model is a simplified version of the Beeler Reuter (BR) [27] model. It comprises three non-linear ODEs. The three variables in the model are the trans-membrane potential (u) and two gating variables (v and w).

The Fitzhugh Nagumo (FHN) [10] model is a simplified version of the Hodgkin-Huxley (HH) model. It comprises two variables,i.e., the excitation variable, v and the recovery variable, u. Additionally, the FHN model introduces a stimulus current (I).

The Fabbri model [11] represents the human sinoatrial node (SAN) cell, commonly known as the pacemaker cell. This model comprises 33 ordinary differential equations (ODEs), with the first ODE describing the AP of the SAN cell. It visualizes and tabulates the relationship between *I*_*f*_ block and the APD variation, which the proposed FM model aims to replicate.

The O’Hara [29] model represents a human ventricular cell. This model comprises 49 ODEs. Jacob et al. [28] used this cell model to create a state controller for the RM. It was utilized to create a 1-D tissue using MATLAB. Using the FM model, 1-D and 2-D tissue models are created by reconstructing the O’Hara [29] model’s action potential duration (APD).

The APDs of these models require simultaneous ODEs. Solvers that take more computational space are required to implement these models on an FPGA. It can lead to practical limitations in generating many cells in real-time. The proposed FM model also aims to overcome these issues.

### Numerical Implementations and Simulations

The concept of the proposed model was simulated in SIMULINK R2022b. The test signals in the reference waveshape section were done using MATLAB R2022b. Later simulations of the reconstructions of the detailed models like Beeler-Reuter (BR), Fenton Karma (FK), FitzHugh Nagumo, Fabbri et al. [11]. The O’Hara [29] cell models were also done on MATLAB R2022b software in the later section. The simulations were conducted on a Windows 10 PC with an Intel Core i7 3 GHz processor. It specifically focuses on determining the duration of the APDs. After this stage, the novel FM model was implemented.

## Results

In this section, we present our findings on 5 different biophysical models. We apply the FM methodology to 4 biophysical models to validate the applicability. Afterward, we show some dynamic properties of the FM cell model, which facilitates us re-emulating the tissue models in 1-D and 2-D. The tissue models capture some dynamic properties using the state controller. A more detailed analysis of other dynamic properties like restitution and refractory period must be addressed.

### Reconstruction of Frequency Modulation factor (Δ_*w*_) plots for Biophysical Models

#### Beeler Reuter (BR) model

Fig. 4(A) depicts the Δ_*w*_ waveshape extracted from BR model. The reconstruction of this plot utilizes 23 breakpoints for a dataset of 40000 values using linear polynomials. Post the optimization method, 22 linear polynomial segments were obtained (see Table.3). Fig. 4(A) illustrates the Δ_*w*_ waveshape and its reconstruction for the BR model.

**Table 3.**
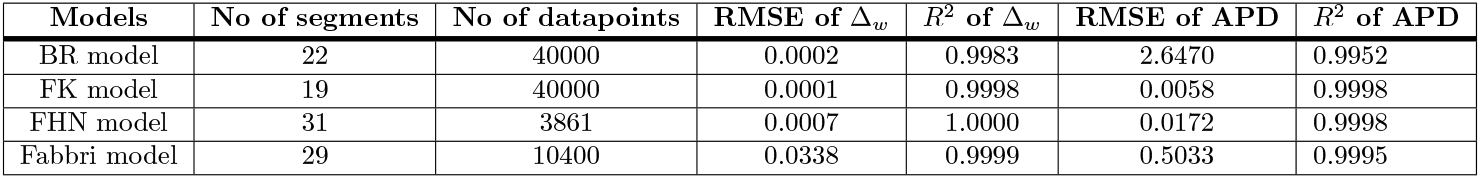
RMSE and *R*^2^ values of the Δ_*w*_ and APD reconstructions using the optimization method for the cell models.

**Fig 4.**
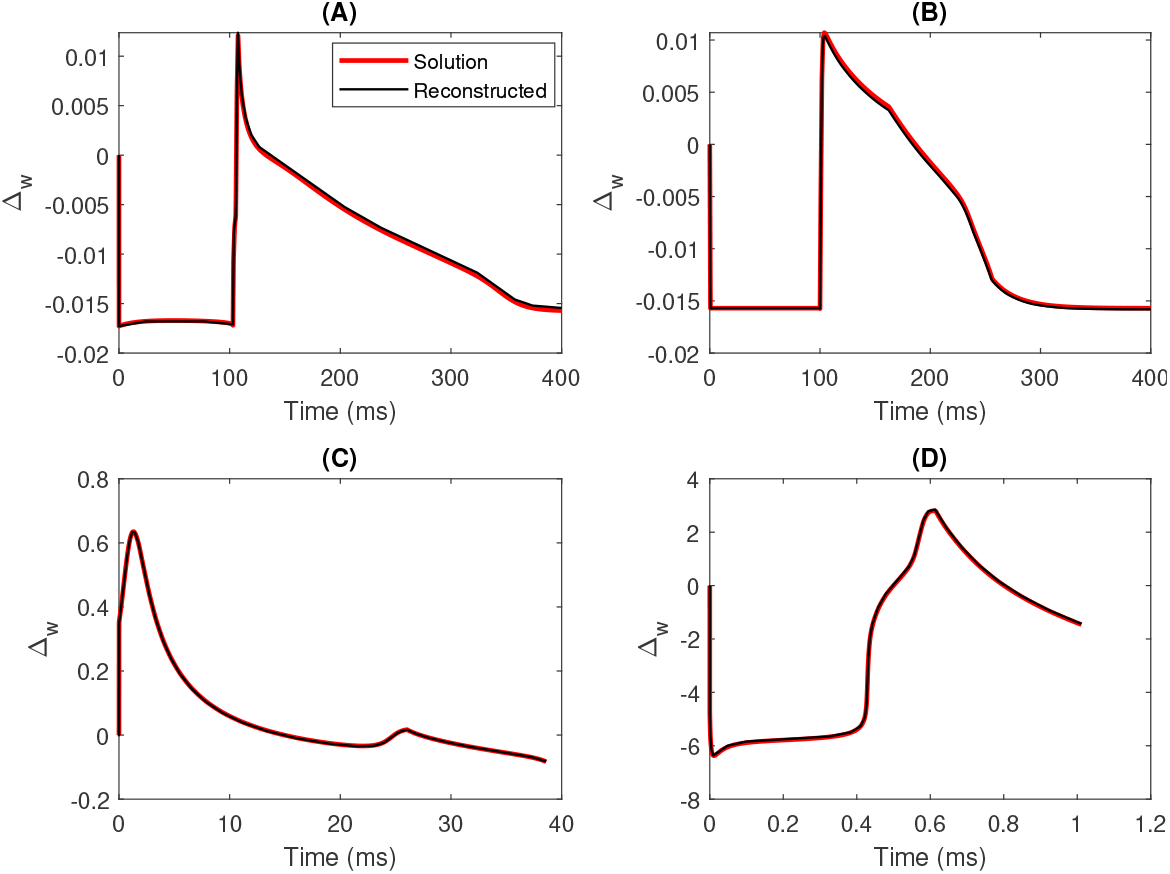
Plots of Δ_*w*_ and its corresponding reconstruction using the optimization method for (A) Beeler-Reuter(BR) [27] model (B) Fenton Karma (FK) [9] model (C) FitzHugh Nagumo (FHN) model and (D) Fabbri [11] model.

#### Fenton Karma (FK) model

Fig. 4(B) shows the Δ_*w*_ waveshape and its corresponding reconstructions. The reconstruction involves 20 breakpoints for a dataset of 40000 values, resulting in 19 linear piecewise polynomials after optimization (refer to Table.3).

#### FitzHugh Nagumo (FHN) model

The FHN model shows the APD of a neuronal cell. Hence, the Δ_*w*_ wave shape for the FHN model (Fig. 4(C)) differs in a big way when compared to BR and FK models. Thrity-two breakpoints were assumed for a dataset of 3861 values. Post-optimization, 31 linear segments were obtained (Table.3). Fig. 4(C) also includes the reconstruction of the Δ_*w*_ plot incorporating the mentioned parameters.

#### Fabbri model

The Fabbri [11] model emulates a pacemaker called the SAN cell. Hence, this model’s Δ_*w*_ waveform varies from the other three cell models. For data of 10400 values, 29 segments were required to attain an optimized reconstruction of the Δ_*w*_ waveshape (Table.3). Fig. 4(D) portrays the reconstruction utilizing the optimized linear polynomials.

Table.3 tabulates the Root Mean Square Error (RMSE) and *R*^2^ error values. The *R*^2^ values are consistently above 0.99 across all the models. However, the BR model quantifies a relatively higher RMSE than other reconstructions. For the FK and FHN models, the RMSE falls below 0.0003.

### Reconstruction of the Action Potential Duration (APD) using Frequency Modulation factor (Δ_*w*_) plots

Upon successfully reconstructing the Δ_*w*_ plots for the mentioned biophysical models, assessing their ability to accurately reconstruct these cell models’ Action Potential Duration (APD) becomes crucial. It is pivotal to observe that the APDs and Δ_*w*_ waveform contain an equal number of data points, as Δ_*w*_ is derived from the APDs of the cell models. Theoretical Background explains Eq.5, which incorporates the frequency modulating factor, Δ_*w*_, with the Action Potential Duration’s (APD’s) frequency, *ω*.

#### Beeler Reuter (BR) model

Fig. 5(A) presents the reconstructed APD of the BR model from the corresponding Δ_*w*_ reconstruction. Despite the slight quantitative offset observed after reaching the peak value, the APD closely resembles the solution.

**Fig 5.**
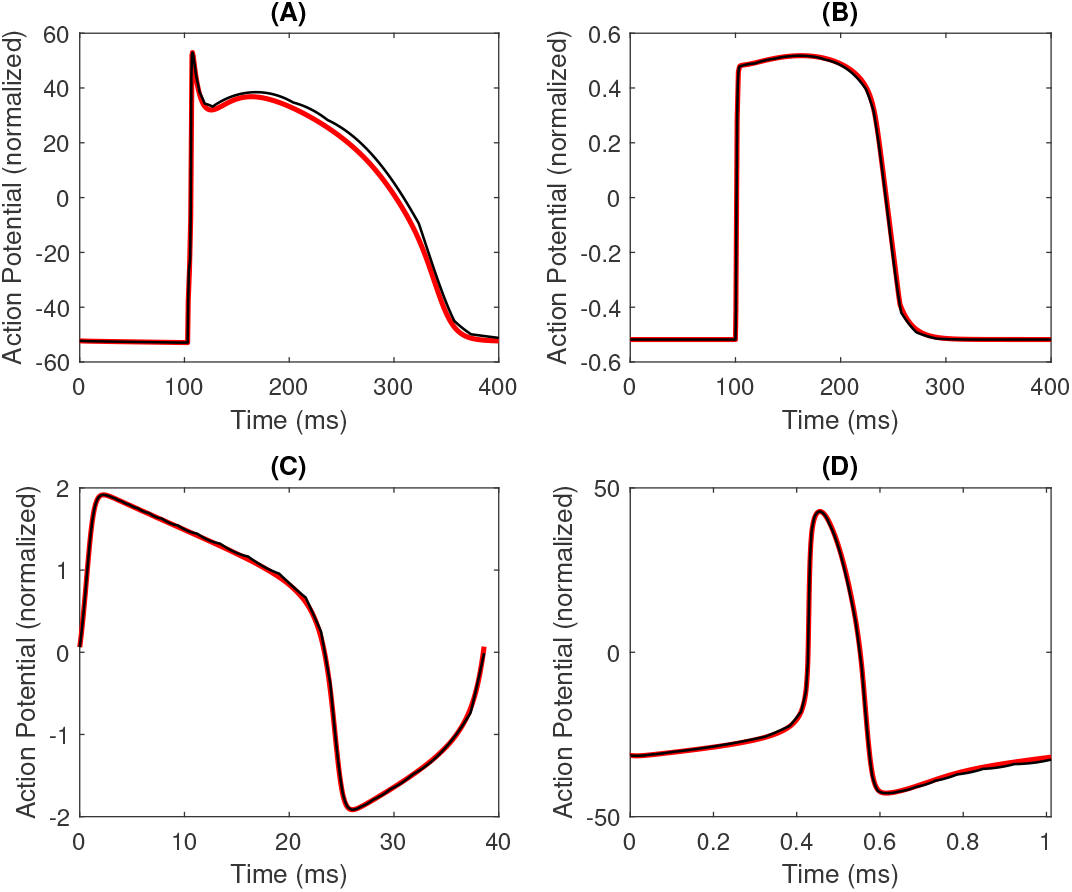
APD waveshape and its corresponding reconstruction for (A) Beeler-Reuter(BR) model (B) Fenton Karma (FK) model (C) FitzHugh Nagumo (FHN) model and (D) Fabbri [11] model.

#### Fenton Karma (FK) model

Fig. 5(B) showcases the reconstruction of the APD from its Δ_*w*_ plot illustrated in Fig. 4(B). A remarkable similarity is observed when the reconstructed APD is compared to the APD of the detailed model. Notably, the number of data points for BR and FK models is 40000.

#### FitzHugh Nagumo (FHN) model

The reconstructed APD of the FHN model is shown in 5(C) which is generated using a 31-segment linear polynomial. The *R*^2^ is found to be more than 0.999, and the RMSE is found to be approximately 0.0172.

#### Fabbri model

The reconstructed APD of the Fabbri [11] model, representing a SAN or the pacemaker cell, is shown in Fig. 5(D). It has *R*^2^ of more than 0.999 and portrays a very low RMSE of approximately 0.5.

Interestingly, despite having similar data points, the BR model replication exhibits higher RMSE than the FK model. The FK model portrays the lowest RMSE (see Table.3) among all the APD reconstructions done. All the cell model APD profiles exhibit a *R*^2^ of above 0.99.

This investigation sheds light on how bankable the FM model is for reconstructing APDs of various cell models. Moreover, the tabulations and illustrations emphasize how effectively the FM model captures the essential characteristics of the original APD profiles.

### Parameterization of the novel FM model

A similar methodology as the RM model [2] incorporating the electrophysiology of ionic current perturbation is done. This study focuses on the simulation of the Fabbri model [11] of the human SAN cell by varying the maximum funny current block, *I*_*f*_, from 0 to 1. Eleven different APD values corresponding to equally spaced *I*_*f*_ values ranging from 0 to 1 are tabulated to investigate the relationship between the APD and the *I*_*f*_ blockade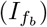.

For recreating the Δ_*w*_ waveshapes of the Fabbri [11] model using this novel methodology, we take 30 breakpoints, which allow us to formulate 29 linear segments with coefficients A1, A2,…., A29. Fig. 6 illustrates the profiles of each polynomial value when subjected to varying 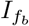 in increments of 0.1. Astonishingly, the trajectories of these polynomial values are linear. Hence, an attempt is made to derive closed-form linear equations that describe the alterations of the polynomial values and the frequency of the dataset, *ω*, for varying 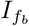. These equations enable the reconstruction of the APDs for an arbitrary 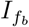.

**Fig 6.**
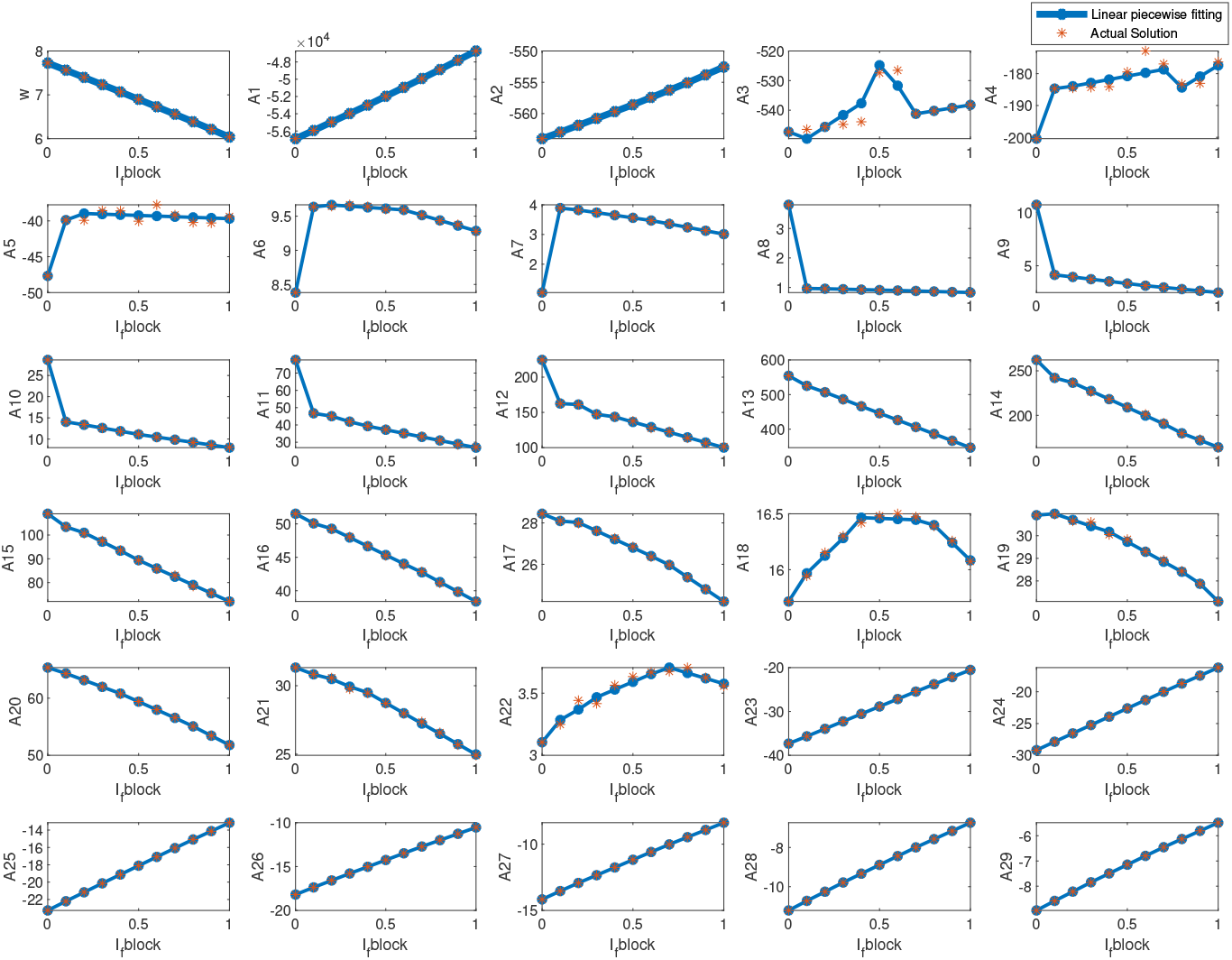
Minimum piecewise linear fits for the polynomial values of Δ_*w*_ of the sinoatrial node cell at different levels of *I*_*f*_ blockade. *ω* is the frequency of the APDs replicated. Δ_*w*_ is divided into 29 different coefficients comprising A1,A2, ……, A29.

Table.4 defines the complexity levels of the fit-type equations. PL3, PL4 and PL5 represent the number of linear piecewise equations required for linear piecewise fitting. PL3 uses 3 linear segments for piecewise fitting. PL4 and PL5 utilize 4 and 5 linear segments for linear piecewise fitting. We use optimization method [19] [20] [21] [22] in Fig. 3(B) to get the location of the breakpoints for a linear piecewise polynomial. The SSE and RMSE enable us to assess the effectiveness and accuracy of the suitable fit types across various 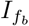. By extracting the fit-type equations for frequency *ω* and 29 coefficients of Δ_*w*_(t) A1, A2, …, A29, we calculate the coefficient of determination (*R*^2^) and the Root Mean Square Error (RMSE) for the reconstructed Δ_*w*_ plots, which corresponds to *I*_*f*_ blockade 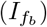 values varying from 0 to 1 in increments of 0.1. For the frequency parameter *ω*, the suitable type used was similar to that of Sehgal et al. [2]. Most coefficients were successfully fitted using the PL4 fit type (refer to Table.5). A5 was fitted using PL3, while A11 and A12 were fitted using PL5. For the fit-type PL4, *ω* has the lowest SSE compared to the other coefficients, followed by A8, A29, A28, A7 and A27 (refer Table.5). However, for PL4, the RMSE is observed to be the lowest for A8, followed by *ω*, A29, A28 and A27. A24 and A26 have relatively the same RMSE. A2 and A6 have the same RMSE. However, the SSE for A2 and A6 are slightly different. A1 had the highest RMSE and SSE when PL4 was applied. However, Fig. 6 illustrates that the fit-type is sufficiently accurate to replicate the values of A1 for varying 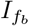 ranging from 0 to 1. A3 and A4 have relatively the same RMSE. However, the SSE for A3 is lesser than that of A4. Interestingly, A13 and A14 have relatively similar RMSE and SSE. A5 utilizes the PL3 fit type and has an RMSE of 0.7181 and SSE of 4.2550. For the PL5 fit-type, A11 exhibits low RMSE and SSE values of approximately 0.15 and 0.24, and A12 exhibits a higher RMSE and SSE of almost 0.86 and 8.2, respectively. Despite the RMSE and SSE discrepancies, we continue to endorse the following fit-types for the coefficients.

**Table 4.** General expression of various fitting functions and the SSE and RMSE for different polynomial values.

| Fit-type | Equation |
| --- | --- |
| PL3 | $\Delta_w(I_{f_b}) = \begin{cases} p_{10} + p_{11} \cdot I_{f_b}, & \text{if } 0 \leq I_{f_b} < r_1 \\ p_{20} + p_{21} \cdot I_{f_b}, & \text{if } r_1 \leq I_{f_b} < r_2 \\ p_{30} + p_{31} \cdot I_{f_b}, & \text{if } r_2 \leq I_{f_b} \leq r_3 \end{cases}$ |
| PL4 | $\Delta_w(I_{f_b}) = \begin{cases} p_{10} + p_{11} \cdot I_{f_b}, & \text{if } 0 \leq I_{f_b} < r_1 \\ p_{20} + p_{21} \cdot I_{f_b}, & \text{if } r_1 \leq I_{f_b} < r_2 \\ p_{30} + p_{31} \cdot I_{f_b}, & \text{if } r_2 \leq I_{f_b} < r_3 \\ p_{40} + p_{41} \cdot I_{f_b}, & \text{if } r_3 \leq I_{f_b} \leq r_4 \end{cases}$ |
| PL5 | $\Delta_w(I_{f_b}) = \begin{cases} p_{10} + p_{11} \cdot I_{f_b}, & \text{if } 0 \leq I_{f_b} < r_1 \\ p_{20} + p_{21} \cdot I_{f_b}, & \text{if } r_1 \leq I_{f_b} < r_2 \\ p_{30} + p_{31} \cdot I_{f_b}, & \text{if } r_2 \leq I_{f_b} < r_3 \\ p_{40} + p_{41} \cdot I_{f_b}, & \text{if } r_3 \leq I_{f_b} < r_4 \\ p_{50} + p_{51} \cdot I_{f_b}, & \text{if } r_4 \leq I_{f_b} \leq r_5 \end{cases}$ |

**Table 5.** RMSE and SSE of each polynomial value post finding the desired fit type.

| Coefficients | RMSE | SSE | Fit-type |
| --- | --- | --- | --- |
| $\omega$ | 0.0007 | $8.39 \times 10^{-7}$ | PL4 |
| A1 | 7.3047 | 93.0648 | PL4 |
| A2 | 0.0111 | $3.46 \times 10^{-4}$ | PL4 |
| A3 | 2.9013 | 0.0017 | PL4 |
| A4 | 2.4492 | 34.3944 | PL3 |
| A5 | 0.7181 | 4.2250 | PL4 |
| A6 | 0.0111 | $3.69 \times 10^{-4}$ | PL4 |
| A7 | 0.0060 | $1.81 \times 10^{-4}$ | PL4 |
| A8 | $3.68 \times 10^{-4}$ | $1.05 \times 10^{-6}$ | PL4 |
| A9 | 0.0114 | $4.60 \times 10^{-4}$ | PL4 |
| A10 | 0.0533 | 0.0132 | PL4 |
| A11 | 0.1493 | 0.2432 | PL5 |
| A12 | 0.8639 | 8.2047 | PL5 |
| A13 | 0.7764 | 2.1277 | PL4 |
| A14 | 0.7149 | 1.9206 | PL4 |
| A15 | 0.2735 | 0.5667 | PL4 |
| A16 | 0.0868 | 0.0593 | PL4 |
| A17 | 0.0285 | 0.0053 | PL4 |
| A18 | 0.0268 | $9.94 \times 10^{-4}$ | PL3 |
| A19 | 0.0772 | 0.0457 | PL4 |
| A20 | 0.0328 | 0.0013 | PL4 |
| A21 | 0.0770 | 0.0294 | PL4 |
| A22 | 0.0375 | $9.92 \times 10^{-4}$ | PL4 |
| A23 | 0.0077 | $2.69 \times 10^{-4}$ | PL4 |
| A24 | 0.0081 | $2.07 \times 10^{-4}$ | PL4 |
| A25 | 0.0095 | $3.05 \times 10^{-4}$ | PL4 |
| A26 | 0.0080 | $3.09 \times 10^{-4}$ | PL4 |
| A27 | 0.0046 | $1.03 \times 10^{-4}$ | PL4 |
| A28 | 0.0042 | $5.16 \times 10^{-5}$ | PL4 |
| A29 | 0.0035 | $4.56 \times 10^{-5}$ | PL4 |

Table.6 tabulates RMSE and *R*^2^ of the reconstructed APDs utilizing the Δ_*w*_ waveshape generated from the fit-type equations to the original APD solution. The RMSE values for the reconstructed Δ_*w*_ vary from approximately 0.016 to 0.05. The highest *R*^2^ for the reconstructed Δ_*w*_ is found for 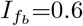 where *R*^2^=1. The RMSE at 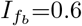 is also the lowest. RMSE of reconstructed APDs vary approximately 0.2 to 0.7. The highest *R*^2^ for the reconstructed APD is found for 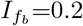 and 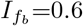 where *R*^2^=0.9999. However, the RMSE for 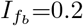 is relatively higher than that of 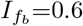. The RMSE is highest, and the *R*^2^ is the lowest for 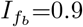 where RMSE=0.7258 and *R*^2^=0.9989. The *R*^2^ for the reconstructed APD varies between 0.9989 to 0.9999. This tabulation showcases that these fit-type equations can effectively replicate the Δ_*w*_ and the AP characteristics when the 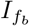 is altered in increments of 0.1 from 0 to 1.

**Table 6.**
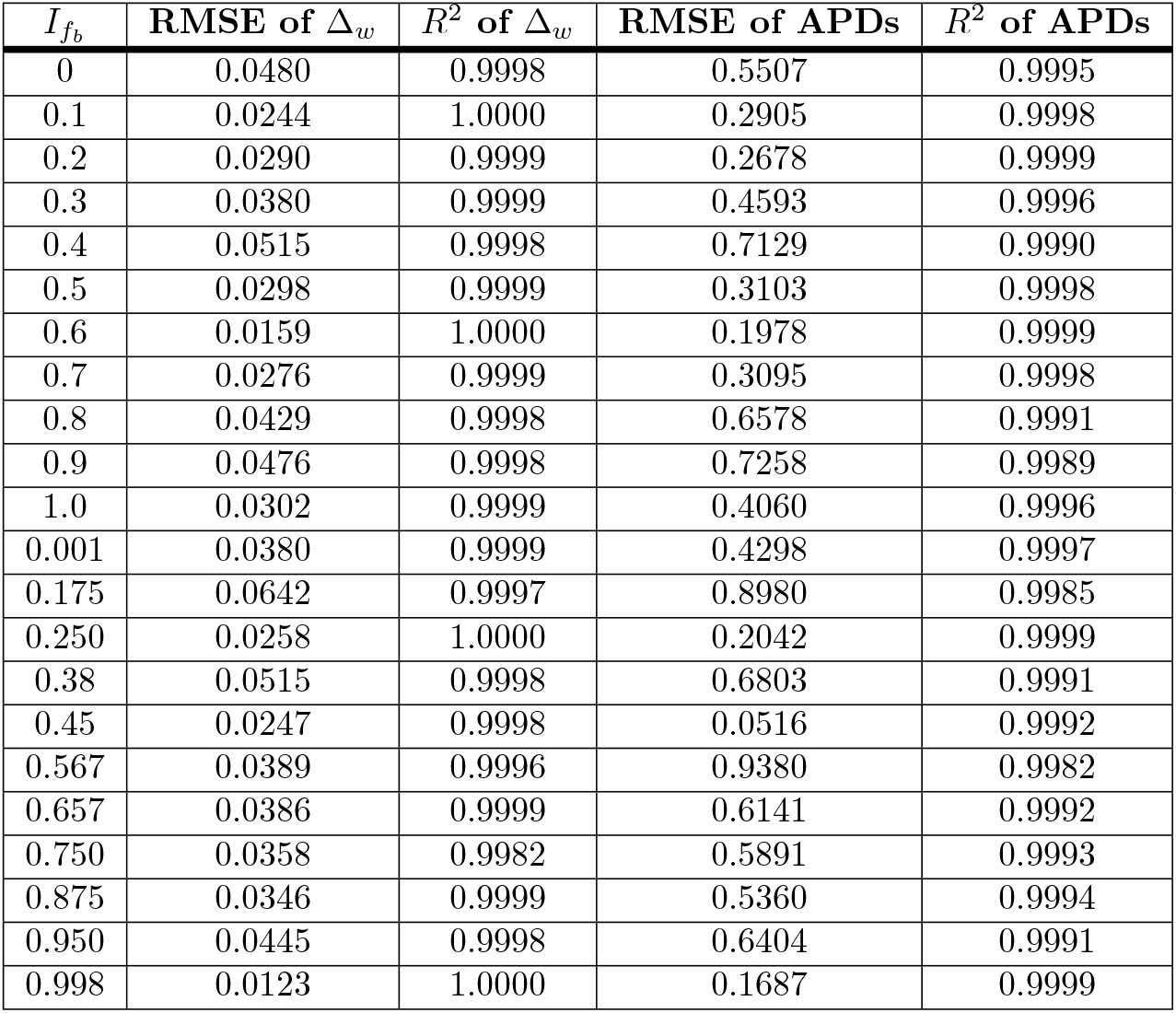
RMSE and *R*^2^ values of the Δ_*w*_ and APD reconstructions using the fit-type equations for different values of *I*_*f*_ blockade.

For evaluating how consistent reconstructed Δ_*w*_ and APD plots are when assigning random values to the *I*_*f*_ blockade 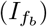, a similar assessment was tabulated to Table.6. It also demonstrates the performance of linear fit-type equations when random values of 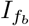 from 0 to 1 are applied. The various values of 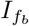are 0.001,0.175,0.25,0.38 and others. The *R*^2^ reveal that the linear fit-type equations consistently reconstruct the Δ_*w*_ waveshape for all the attempted random values of 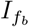, as all of them are above 0.998 (refer Table.6). RMSE of the Δ_*w*_ replication ranges from 0.0123 to 0.0642. Notably, a surge in *R*^2^ corresponds to a decrease in RMSE, reinforcing the effectiveness of the linear fit-type equations in capturing the Δ_*w*_ characteristics for various *I*_*f*_ blockade, 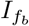. Table.6 also quantitatively analyzes how the random 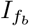 values from 0 to 1 can reconstruct the APD waveshapes. Despite a higher RMSE for the APD waveshape replication, the *R*^2^ is consistently above 0.998. The highest RMSE is at 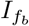 =0.567, corresponding to the lowest the lowest *R*^2^,i.e., 0.9982. The RMSE for the APD reconstruction is also the lowest at 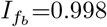, corresponding to the highest *R*^2^, i.e., 0.9999. To sum up, the linear fit-type equations successfully replicate the Δ_*w*_ and AP characteristics of the SAN call not only for 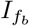 values in increments of 0.1 but also for arbitrary 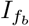 values between 0 to 1.

Fig. 7 (A) and (B) visually illustrate the alterations in the Δ_*w*_ waveshape for 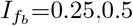 and 0.75. In addition to this, Fig. 7 (C) and (D) depict how the AP characteristics vary when the *I*_*f*_ blockade, 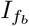varied at 0.25,0.5, and 0.75. These figures portray how the reconstructed waveshapes closely resemble the waveshapes of the detailed model when subjected to a particular *I*_*f*_ blockade, 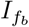. Hence, we have conclusive evidence that the FM model can alter the AP characteristics of a SAN cell when there is an alteration in the values of *I*_*f*_ blockade, 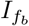.

**Fig 7.**
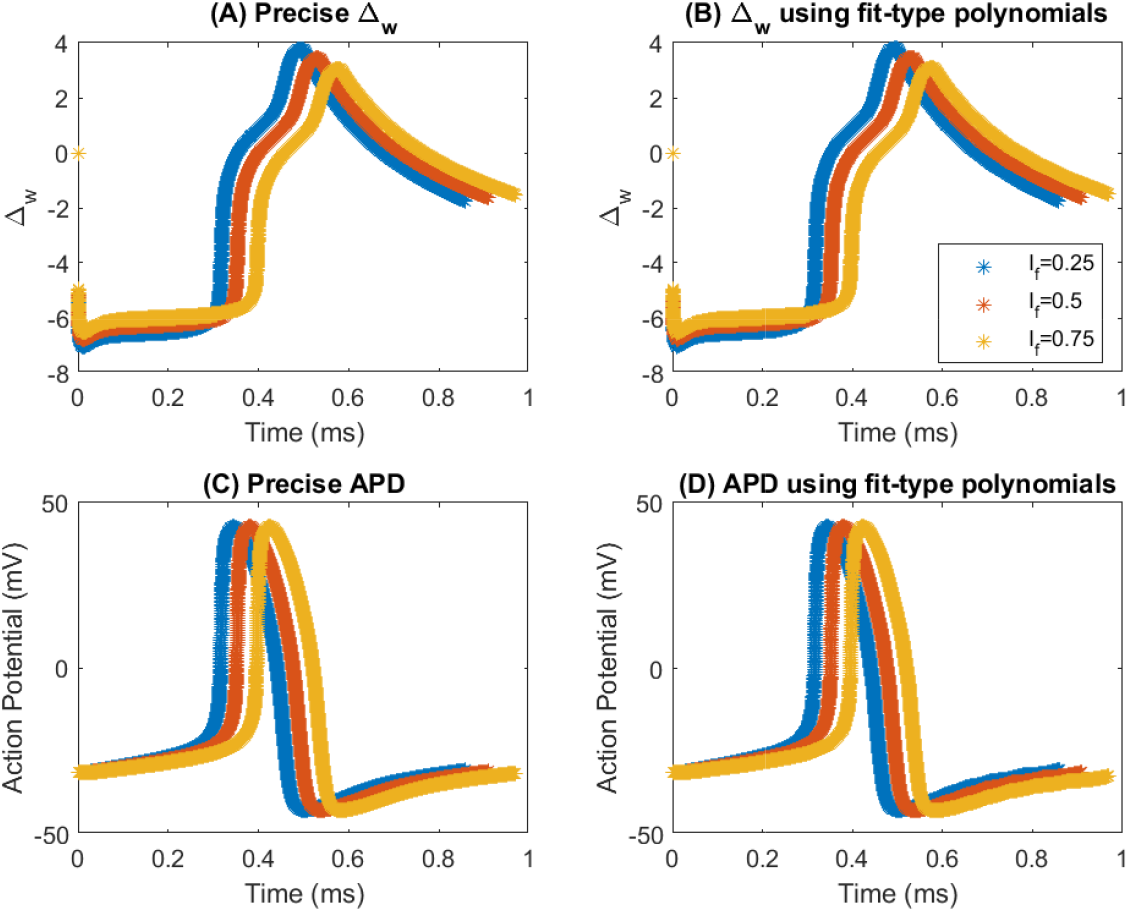
Variation of Δ_*w*_ waveshape and APD when 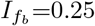,0.50 and 0.75 (A) Precise Δ_*w*_ (B) Δ_*w*_ using fit-type polynomials (C) Precise APD (D) APD using fit-type polynomials.

The investigation in this subsection affirms that the FM model can accurately reconstruct the APD of the human SAN cell for any 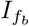 ranging from 0 to 1.

### Creating dynamic behavior to the FM cell model

This part of the work discusses creating more complexity and capability for the reconstructed AP characteristics using the FM methodology. Sehgal et al. [1] discuss the state controller for reconstructing the O’Hara [29] model. Jacob et al. [3] utilized it to create a 1-D tissue. Hence, we need to reconstruct the APD of the O’Hara [29] cell model using the FM model. Fig. 8 (A) and (B) showcase the replication of the O’Hara [29] cell model. To reconstruct this model’s Δ_*w*_ profile, we used 21 segments, i.e., 22 breakpoints for 50001 data points. The replication of the Δ_*w*_ profile is illustrated in Fig. 8 (A). Like any other cell model’s reconstruction, we calculated the RMSE and *R*^2^ of the reconstructed Δ_*w*_ waveshape. The RMSE of the reconstructed Δ_*w*_ profile was as low as 0.0002, and the *R*^2^ was approximately 0.9993. The APD replication is found to have low RMSE, corresponding to the high *R*^2^ value.

**Fig 8.**
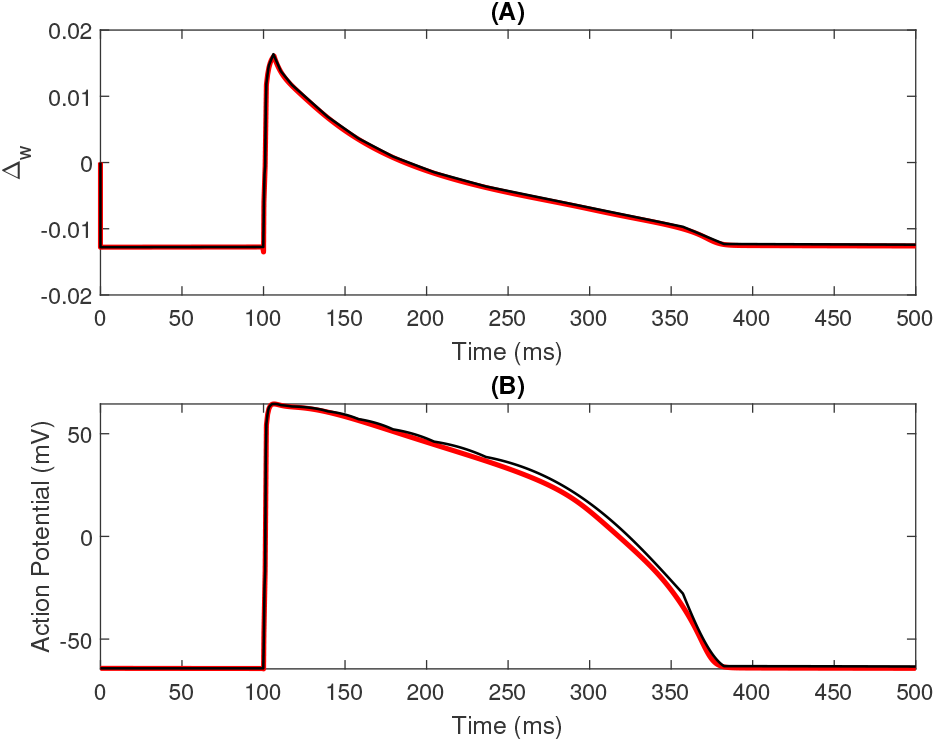
Reconstruction of the (A) the Δ_*w*_ profile and (B) the APD replication of the O’Hara [29] cell model.

Sehgal et al. [1] had three states in their state controller comprising *S*_1_: resting state, *S*_2_: Upstroke, and *S*_3_: Repolarization states. These states present in the state controller of the RM are bounded by a certain set of conditions. Fig. 9 shows the point-to-point variations of the APD reconstruction of RM with and without state controller along with the replication using the FM model. Table.7 depicts the RMSE and *R*^2^ of the O’Hara [29] APD replication using the three techniques. It can be observed that the RMSE of the RM cell without a state controller is significantly high, i.e., 5.3603. This metric affects the *R*^2^ and is the lowest among the three. On utilizing RM with state controller, the RMSE dwindles significantly from 5.3603 to 2.1849, which also shoots up the *R*^2^. The FM model replication’s RMSE and *R*^2^ values are almost similar to that of the RM with the state controller.

**Table 7.** The RMSE and *R*^2^ values of O’Hara [29] model reconstructions using RM with/without the state controller and the FM model.

| Method | RMSE | $R^2$ |
| --- | --- | --- |
| RM without state controller | 5.3603 | 0.9885 |
| RM with state controller | 2.1849 | 0.9985 |
| FM model | 2.4601 | 0.9976 |

**Fig 9.**
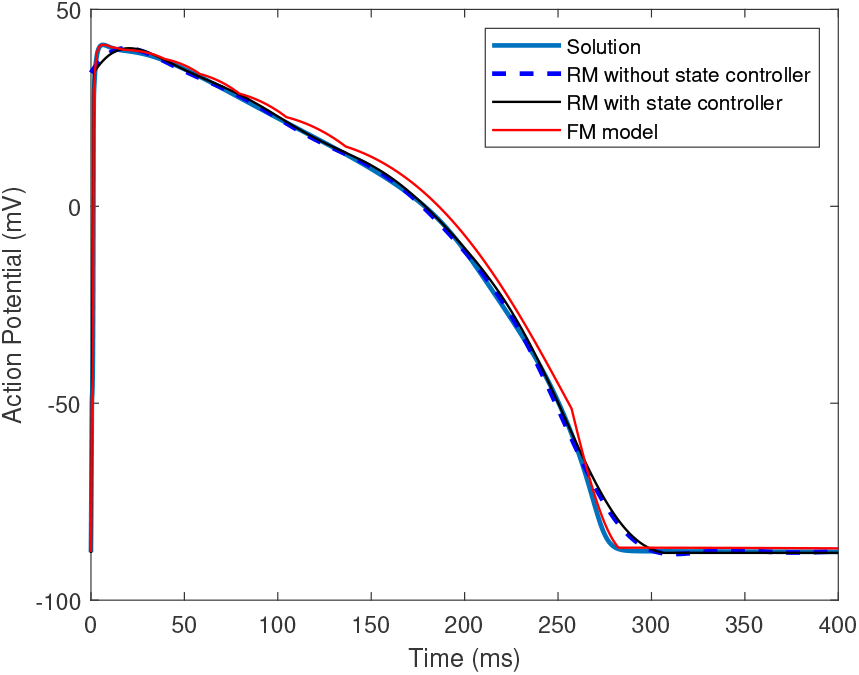
The point-to-point difference in APD reconstruction w.r.t solution for RM without state controller, RM with state controller and FM model.

For the current work, we designed a state controller comprising two states, i.e., State 1:resting state and State 2: APD generation state. Fig. 10 shows the state controller of the FM model. Various constants and variables are required for this state controller. These include the variable, S, threshold voltage, *V*_*th*_, resting voltage, *V*_*rest*_, frequency modulating factor at rest, Δ_*rest*_, frequency modulating factor of the APD, Δ_*w*_, stimulus voltage, *V*_*sti*_ and *V*_*out*_, the voltage of the generated APD.

**Fig 10.**
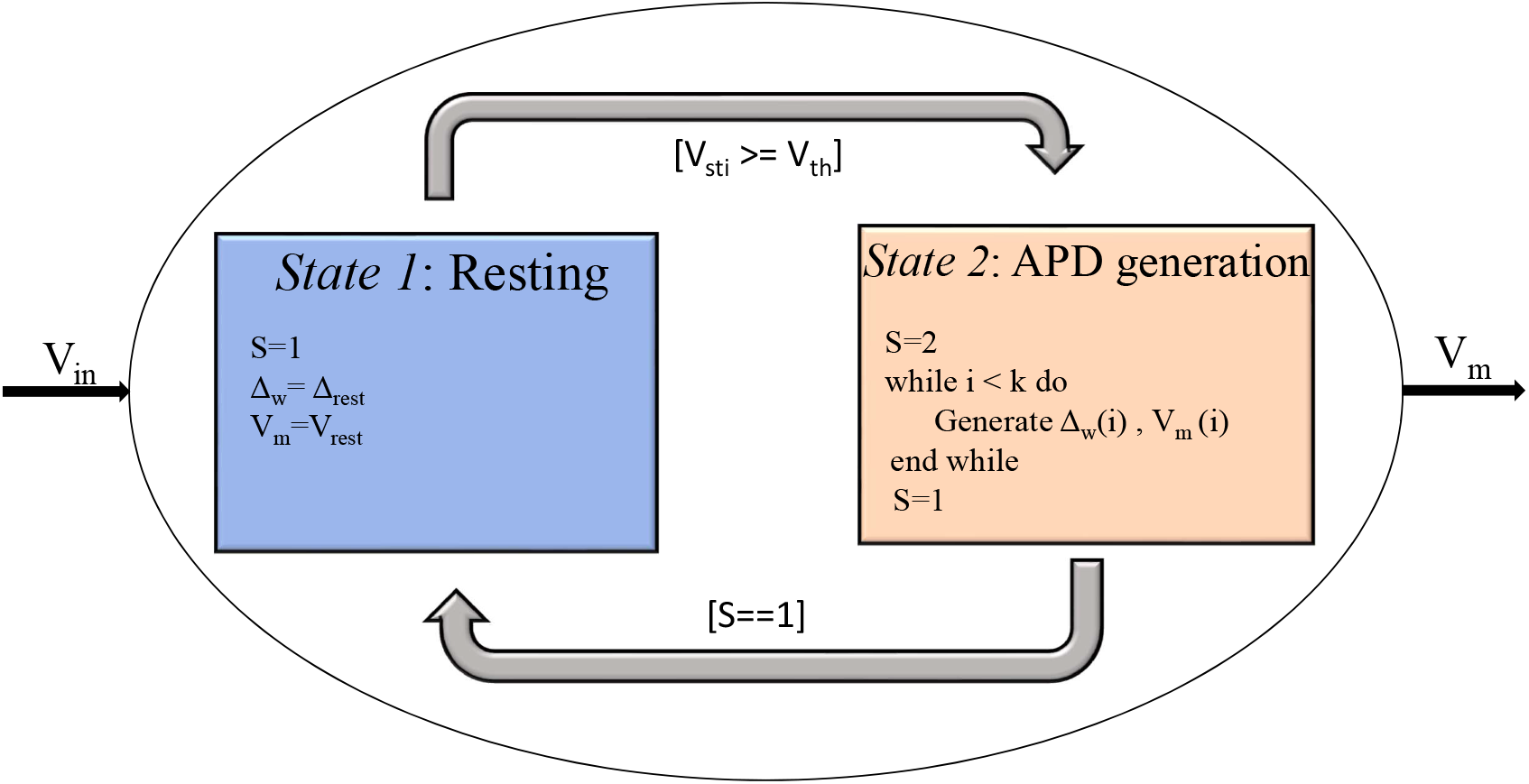
State controller of the FM model to create dynamic behavior for the FM model.

At State 1 (when S==1), the cell remains at rest; hence, the membrane voltage, *V*_*m*_, is at the resting voltage, *V*_*rest*_. The Δ_*w*_ is equivalent to the initial value of the Δ_*w*_ profile generated for the cell model denoted as Δ_*rest*_. The arrival of a stimulus with *V*_*sti*_ *>*= *V*_*th*_ causes the cell to switch to State 2, where S transitions from 1 to 2. In this state, the APD is generated for a defined number of samples, k, after which the value of S becomes 1. When S==1, the cell goes back to the resting state.

The FM model reconstruction of the O’Hara [29] model was done where the upstroke and repolarization state begins at 100 ms. Hence, it is imperative to know whether the state controller allows the cell stimulation at 0 ms. When the stimulus was applied at 0 ms, as depicted in Fig. 11 (a), S begins at 2 rather than 1. Hence, we can infer that the cell reacts to the stimulus at 0 ms. We can observe that S remains at 2 till 300 ms. After 300 ms, we can infer that the stimulus applied to the cell is significantly low, so the cell remains resting where S is 1. In Fig. 11 (b), the stimulus is applied at 200 ms for the cell. From 0 to 200 ms, S was at 1,i.e., the cell was at rest during this period. The state shifts from 1 to 2 after 200 ms, i.e., the cell has entered the APD state. The APD state is found to be ending at 500 ms when the state, S, is altered from 2 to 1. Fig. 11 (c) illustrates a similar waveform and state evolution as that of Fig. 11 (a) and (b) when the stimulus is applied at 350 ms. Similar observations and analysis should be done for the APD reconstructions of the FHN and Fabbri et al. [11] model.

**Fig 11.**
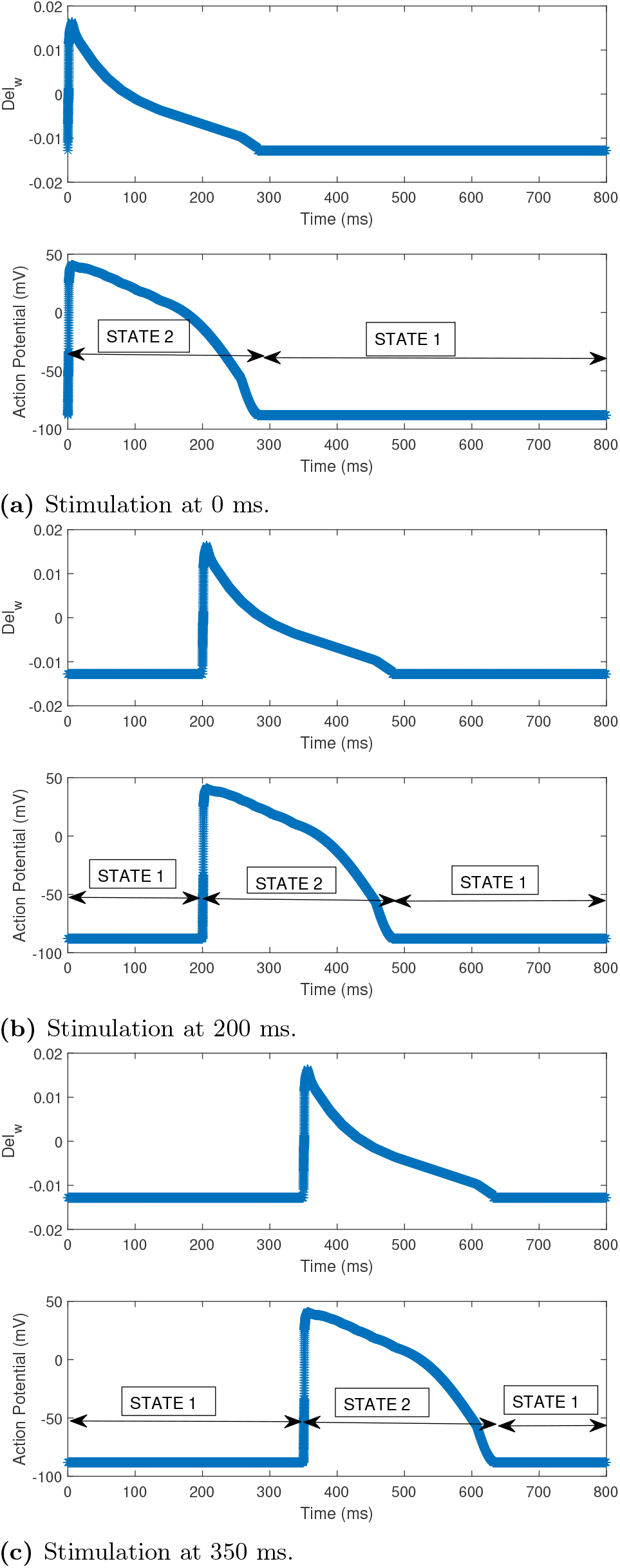
The Δ_*w*_ and APD reconstructions of the O’Hara [29] model along with the alterations in the state value in the FM state controller when the Stimulation is at (a) 0 ms (b) 200 ms and (c) 350 ms.

Since the reconstruction of the FHN model using the FM model began at 0 ms, we do not need to test whether the FM state controller stimulates the AP characteristics at that instant. Hence, we look to execute the state controller for the replicated APD of the FHN model at other time instants like 20 ms, 30 ms and 35 ms. Fig. 12 (a) showcases how the AP characteristics of the reconstructed FHN model propagates at 20 ms. From the time instants between 0 ms to 20 ms, the state value of the state controller remains at 1. After 20 ms, we can observe that S alters from 1 to 2. When the S is at 2, the APD of the FHN model is generated exactly as the reconstructed FHN model using the FM model. When the S is 2, the APD of the FHN model is generated exactly as the reconstructed FHN model as per the FM methodology. The APD propagation ends at 60 ms. Afterward, S alters itself from 2 to 1. On altering the time instant of the stimulus from 20 ms to 30 ms, we get Fig. 12 (b). In this case, S is 1 from 0 ms to 30 ms. After 30 ms, the APD propagates with the reconstructed characteristics of the FHN model. The propagation starts from 30 ms and ends at approximately 70 ms. The S value of the state controller alters from state 1 to 2 in this period. When the time instant is after 70 ms, S alters from 2 to 1, showing that the FM cell is at rest. To further test the state controller with this cell model, we altered the time instant of stimulation from 30 ms to 35 ms (refer Fig. 12 (c)). The FM cell remains at the APD state from 35 ms to approximately 75 ms. Afterward, S alters from 2 to 1, showing that after 75 ms, the FM cell remains resting.

**Fig 12.**
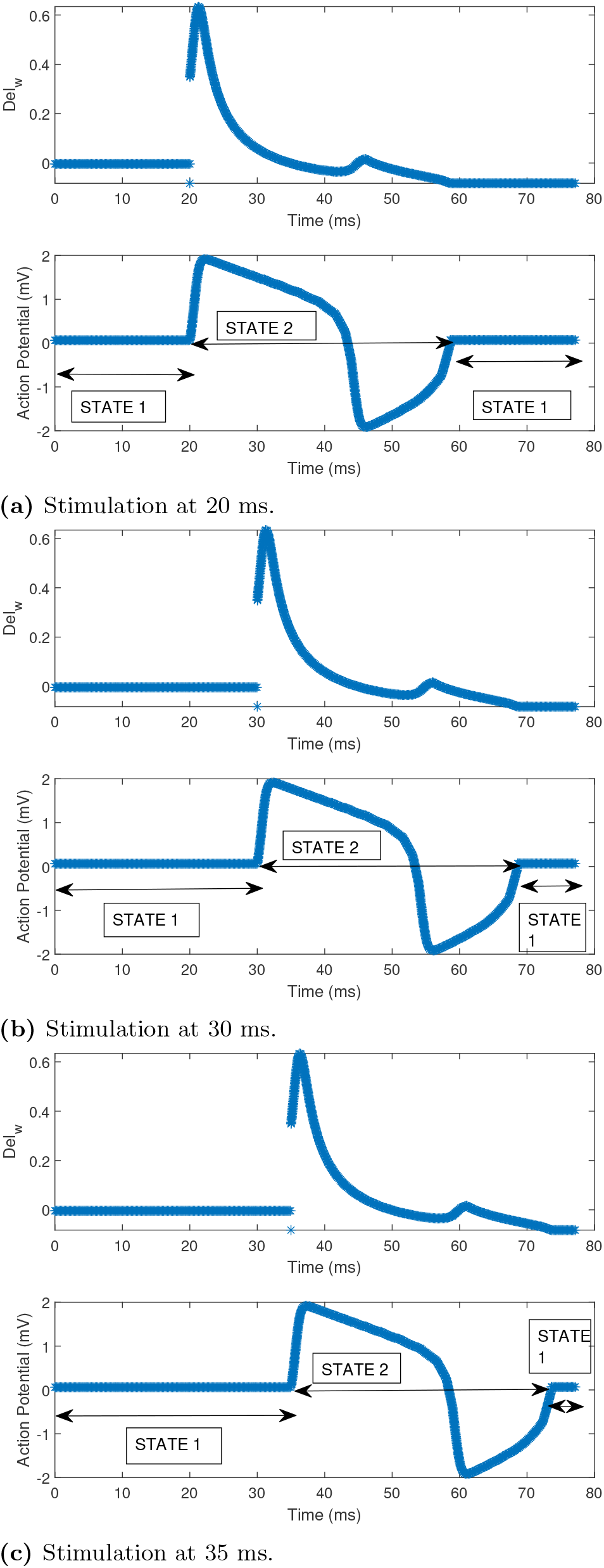
The Δ_*w*_ and APD reconstructions of the FHN model along with the alterations in the state value in the FM state controller when the stimulation is at (a) 20 ms, (b) 30 ms and (c) 35 ms.

Similar observations can be found when we use the Fabbri [11] model APD for the state controller. We have designated time points at 0.5 ms, 0.75 ms, and 0.9 ms for applying the stimulus. These can be found in Fig. 13 (a), (b) and (c). We can observe S altering from 1 to 2 when the stimulus is applied in the earlier instances. After the APD state, S moves from 2 to 1, showing that the FM cell has transitioned from APD to resting.

**Fig 13.**
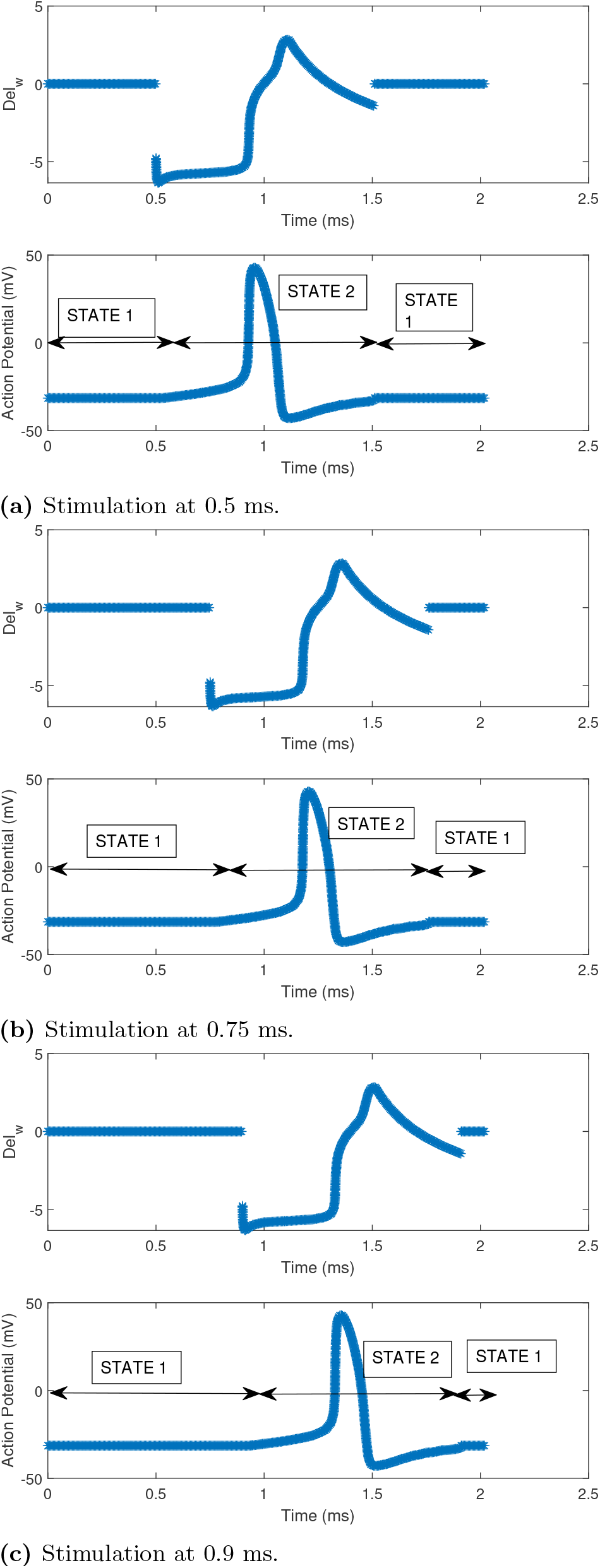
The Δ_*w*_ and APD reconstructions of the Fabrri model along with the alterations in the state value in the FM state controller when the stimulation is at (a) 0.5 ms, (b) 0.75 ms and (c) 0.9 ms.

### Emulation of 1-D FM tissues

Previously, we have implemented a state controller of the FM cell model, which creates complexity and capability for the reconstructed APDs of the cell models such as O’Hara [29], Fabbri et al. [11] and FHN models. Fig. 14 illustrates the design for connecting 1-D cells using the FM model. Each cell comprises the state controller, connected using the diffusion equations discussed in Eq13.

**Fig 14.**
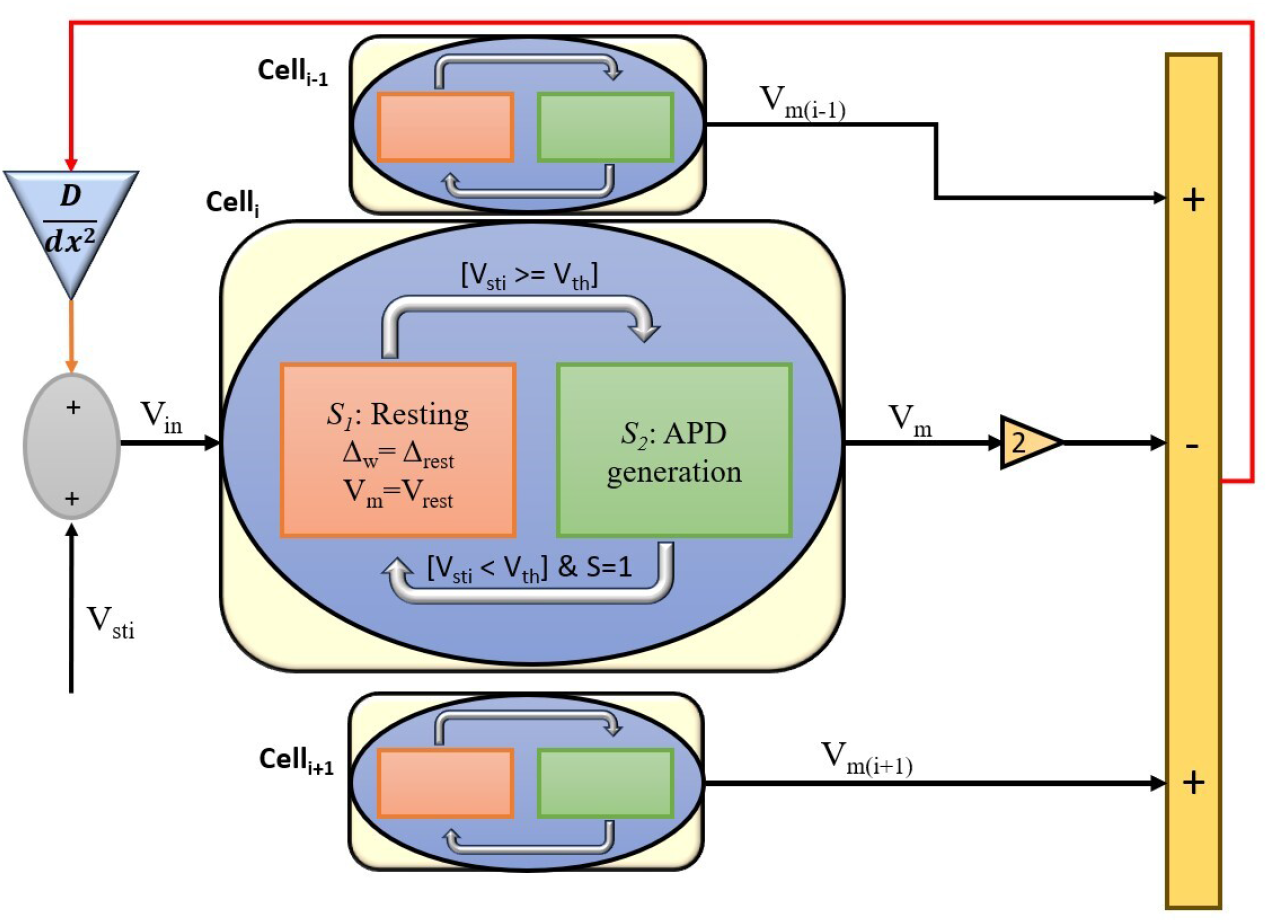
The potential design for connecting FM cells in a 1D tissue. The detailed FM state controller is shown for the *i*^*th*^ cell. D is the diffusion coefficient. *dx* is the distance between the neighboring cells.

Fig. 15 (A), (B), and (C) illustrate the 1-D propagations wavefront for O’Hara [29], FHN, and Fabbri [11] model where the stimulus is applied at cell 1, cell 8, and cell 15. The diffusion constant, D and step time for Fig. 15 (A) is 0.0625 and 0.4 ms. While for Fig. 15 (B), these are 0.00375 and 0.1 ms, and for Fig. 15 (C), these are 0.1052 and 0.02 ms. In Fig. 15 (A) and (B), when the stimulus is applied at cell 1, the wavefront propagates from cell 1 to cell 15. When the stimulus is altered from cell 1 to 8, the wavefront propagates outwardly, i.e., from cell 8 to cell 1 and cell 8 to cell 15. When cell 15 is stimulated, the propagation wavefront evolves from cell 15 to cell 1. However, the evolution of the propagation wavefront of the Fabbri [11] model (refer Fig. 15 (C)) is different from the O’Hara [29] and FHN model. This difference is due to the autorhythmic nature of the Fabbri [11] cell model. In this case, when the stimulus is applied at cell 1, the wavefront propagates from cell 1 to cell 15 at regular intervals. As the stimulus is altered from cell 1 to cell 8, the wavefront evolves outward at regular intervals, i.e., from cell 8 to cell 1 and from cell 8 to cell 15. As the stimulus is exerted on cell 15, the wavefront regularly propagates from cell 15 to cell 1. These illustrations show that the 1-D diffusion equation can propagate autorhythmic cells and those that require excitation.

**Fig 15.**
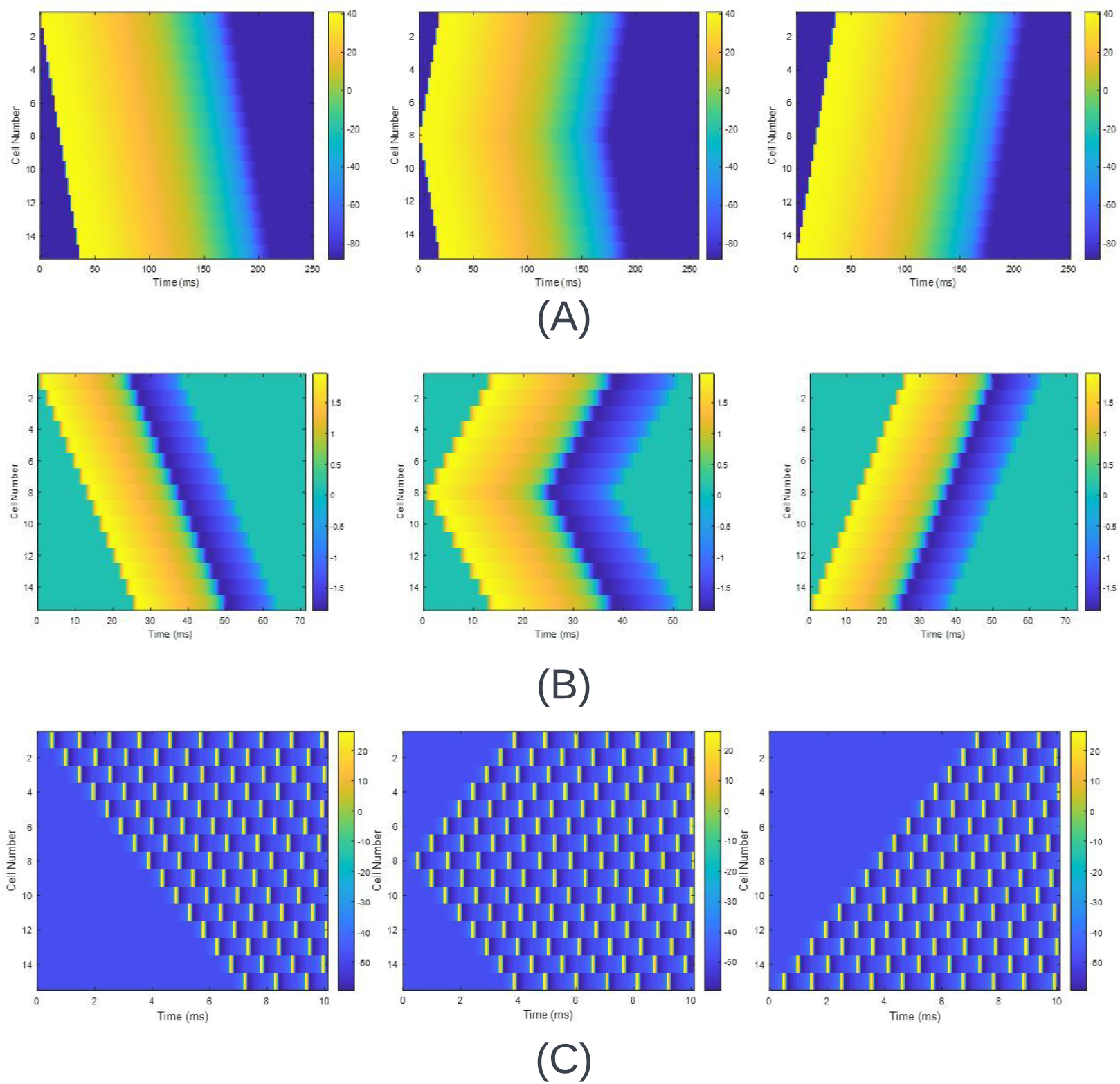
The propagation wavefront of 1-D FM model tissue comprising 15 cells for three different junctures of stimulation for (A) O’Hara [29] model where D=0.0625 for step time of 0.4 ms (B) FHN model where D=0.00375 for step time of 0.1 ms (C) Fabbri [11] model where D=0.1052 for a step time of 0.01 ms.

### Emulation of 2-D FM tissues

Earlier, we showed the wavefront propagation of the 1-D tissue using the FM model. We will observe how a 2-D FM tissue is governed by the diffusion equation in Eq18 for O’Hara [29], FHN, and Fabbri models. The cell assignment in a 2-D tissue is done in a systematic pattern. In a 15×15 2-D tissue, the cell at the top-left corner is cell 1, while the cell at the bottom-right corner is cell *N* ^2^,i.e., 225, and the middle cell can be found using (*N* ^2^ + 1)*/*2, i.e., 113. Initially, we attempted to find the propagation wavefront of a 2-D FM tissue where the stimulus is applied at 4 different junctures at (1) cell 1, (2) cell (*N* ^2^ + 1)*/*2, (3) cell *N* ^2^ and (4) cell 1 to N in the vertical direction for a 7×7,15×15 and 31×31 2-D FM tissue.

Fig. 16 (A), (B), (C) and (D) showcase the propagation wavefront when the stimulus is exerted on a 2-D tissue comprising 15×15 cells. The diffusion constant and step time are 0.0625 and 0.4 ms. When cell 1 is stimulated, the propagation wavefront evolves, as shown in Fig 16 (A). The wavefront propagates outwardly, affecting all the adjacent cells from cell 1 to cell *N* ^2^=225. When cell 1 is found to go to the resting state, we can observe that all the cells within the 2-D tissue gradually go to the resting state. When the stimulus exerted is shifted from cell 1 to cell (*N* ^2^+1)/2, i.e.,113, we get the wavefront to propagate as in Fig 16 (B). The wavefront propagates outwardly in all directions, affecting the surrounding cells within the 2-D tissue. As cell 113 is found to be at resting condition, all the cells are within the 2-D tissue. Later, we look to shift the stimulus from cell (*N* ^2^+1)/2 to cell *N* ^2^, i.e., 225 in this case. The illustrations are displayed in Fig 16 (C). The wavefront propagates opposite to the illustration showcased in Fig 16 (A). The propagation begins from cell 225 to cell 1, diagonally affecting the adjacent cells. As cell 225 goes to rest, all the cells within the 2-D FM tissue gradually do so. Afterward, as shown in Fig 16 (D), we stimulated cells 1 to 15 together in the vertical direction. In this case, the wavefront propagates linearly in the vertical direction. When cells 1 to 15 go to the resting state, we can observe that all the adjacent cells tend to go to the resting state. Similar observations were found for the 7×7 and 31×31 2-D FM tissues.

**Fig 16.**
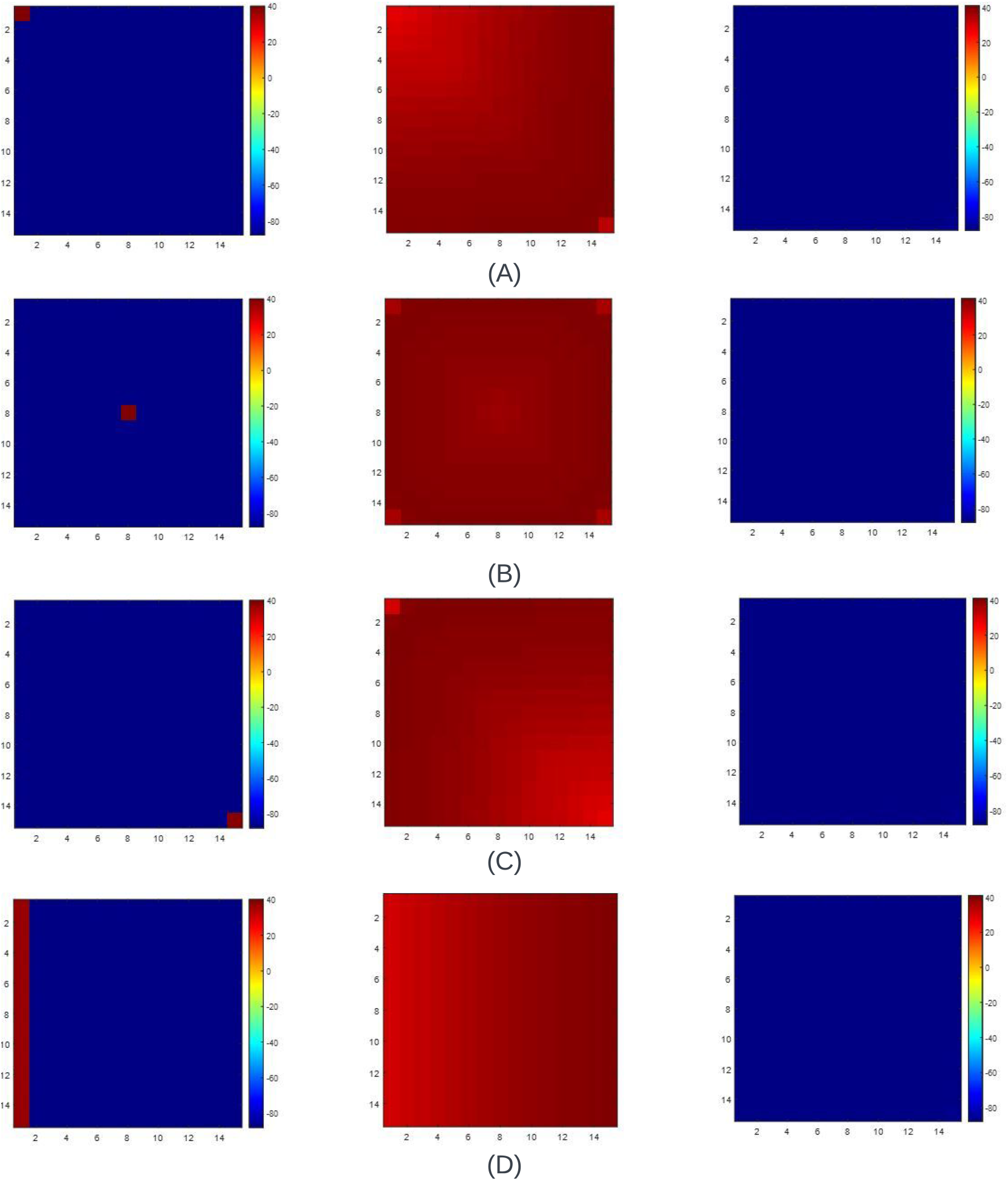
The propagation wavefront of a 15×15 2-D tissue of the O’Hara [29] model when the (A) Cell 1 is stimulated (B) Cell (*N* ^2^+1)/2 is stimulated, i.e., Cell 113 for a 15×15 cell matrix is stimulated (C) Cell NxN, i.e., 225 is stimulated and (D) Multiple cell from Cell 1 to N is stimulated where N=15 is stimulated. The propagation wavefront starts differently for each stimulation, but in the end, all the cells in the 2-D tissue rest when the cells or cells are stimulated to return to rest. The diffusion constant, D, used in all cases is 0.0625, and the step time is 0.4 ms.

Various parameters are required to justify working a 7×7, 15×15 and 31×31 2-D FM tissue. These are

1. Peak time at cell 1,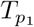: The time at which cell 1 reaches the value of the maximum membrane potential. It is measured in milliseconds (ms).
2. Mid-cell peak time,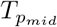 (ms): The time at which the cell (*N* ^2^+1)/2 reaches the maximum membrane potential.
3. Peak time at cell *N* ^2^, *T*_*pNN*_ (ms): The time at which the cell *N* ^2^ reaches the maximum membrane potential.
4. Mid to cell 1 peak time, *T*_*mc*1_ (ms): The absolute difference in peak time at cell 1 to the mid-cell peak time. The equation is depicted in Eq19.

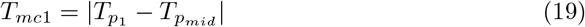
5. Mid to cell NxN peak time, *T*_*mNN*_ (ms): The absolute difference in peak time at cell *N* ^2^ to the mid cell’s peak time. The formulation of this parameter is shown in Eq20.

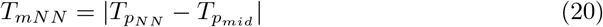
6. Relative adjacent peak time, *T*_*adj*_ (ms): The absolute difference between the peak time of the first stimulated cell and the peak time of the adjacent cells.

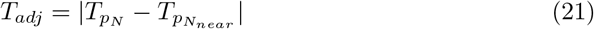

where *T*_*pN*_ is the peak time of the initially stimulated cell and 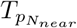 is the peak time of the nearby cell, which is influenced by the stimulation of the initial cell. For example, for a 15×15 2-D tissue, when cell, N=(*N* ^2^ + 1)*/*2=113 is excited, the *T*_*adj*_ can be

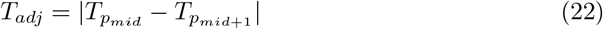

or

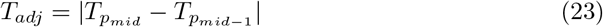

or

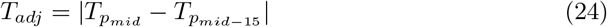

or

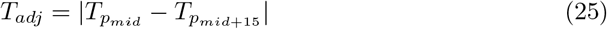

where 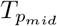 is the peak time at cell (*N* ^2^ + 1)*/*2=113, 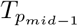 is the peak time at cell ((*N* ^2^ + 1)*/*2)-1=112, 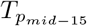 is the peak time at cell ((*N* ^2^ + 1)*/*2)-15=98 and 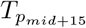 is the peak time at cell ((*N* ^2^ + 1)*/*2)+15=128 for a 15×15 2-D tissue.

For all the Eqs 22, 23, 24 and 25, *T*_*adj*_ should be the same in all the cases. For creating the wavefront propagation of 2-D FM ventricular myocyte, we have found that the minimum value for the diffusion constant, D, equals 0.0625. We calculated the metrics for all conditions discussed using this value and the step time of 0.4 ms as a constant. Table.8 displays these values when stimulated by the cell (*N* ^2^ + 1)*/*2. We can observe that the values of 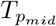 and *T*_*adj*_ are the same, i.e., 7.6 ms and 4.8 ms, respectively, despite the alteration in the size of the 2-D FM tissue. This observation shows that when the cell in the middle is stimulated, the wavefront propagates despite the alteration in size. In all cases, the values of *T*_*p*1_ and *T*_*pNN*_ increase as the size of the 2-D FM tissue expands. Despite the observation of the surge, the individual values of *T*_*p*1_ and *T*_*pNN*_ are similar for a 7×7,15×15 and 31×31 2-D FM tissue depicting that the wavefront propagates outwardly. These properties are also observed when we tabulate the values for *T*_*mc*1_ and *T*_*mNN*_. As the size of the tissue expands from 7×7 to 15×15, the values of *T*_*p*1_, *T*_*pNN*_, *T*_*mc*1_ and *T*_*mNN*_ is found to be doubled. The parameter further doubles as the size expands from 15×15 to 31×31. This analysis shows that the 2-D diffusion equation works for the FM ventricular tissue. It also displays the variations of the parameters when cell 1 is stimulated for a diffusion constant of 0.0625 and a step time of 0.4 ms. In this case, the parameter *T*_*p*1_ is observed to be the same in all cases. Hence, we can infer that we have stimulated cell 1 in all the cases. The values of *T*_*adj*_ are similar despite the alterations in the size of the 2-D FM tissue. In all cases, the values of 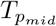 are lesser than that of *T*_*pNN*_, showing proof of how the wavefront propagates when cell 1 is stimulated. The values of *T*_*mc*1_ and *T*_*mNN*_ doubles when the size of the 2-D FM tissue evolves from 7×7 to 15×15. These values tend almost to quintuple as the 2-D FM tissues expand from 7×7 to 31×31. Similar observation can be inferred for 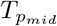 and *T*_*pNN*_. Through this analysis, we can quantitatively comprehend the evolution of the wavefront propagation when cell 1 is stimulated. It also shows the variation of the same metrics when the cell *N* ^2^ is stimulated using the same diffusion constant and step time. In this case, the value of *T*_*pNN*_ is the same, showing that the cell *N* ^2^ is stimulated. The value of *T*_*p*1_ in this case is found to be equal to that of the *T*_*pNN*_ when cell 1 is stimulated. 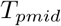 is the same as the values found when cell 1 is stimulated. *T*_*mc*1_ is similar to that of *T*_*mNN*_ when cell 1 is excited. Moreover, *T*_*mNN*_ is equal to *T*_*mc*1_ when cell 1 is excited. This tabulation infers that the wavefront propagation when cell *N* ^2^ is stimulated is opposite to the wavefront propagation when cell 1 is stimulated. When cells 1 to N are stimulated in the vertical direction, we get the parameters shown in Table.8. In this case, the values of *T*_*p*1_ and *T*_*adj*_ are the same across all the 2-D model sizes. *T*_*mc*1_ and *T*_*mNN*_ are the same despite the alteration in the size of the 2-D tissue. The 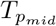 and *T*_*pNN*_ doubles as the tissue size alters from 7×7 to 15×15. The values almost double when the 2-D FM tissue size alters from 15×15 to 31×31. This analysis explains how the values of the respective parameters alter when a set of cells from 1 to N is stimulated.

**Table 8.** The parameters of time for 7×7,15×15 and 31×31 2-D O’Hara tissue when the diffusion constant, D=0.0625 for different junctures of stimulation. The step time is 0.4 ms.

| Stimulation | 2-D model size | $T_{p1}$ (ms) | $T_{pmid}$ (ms) | $T_{pNN}$ (ms) | $T_{mc1}$ (ms) | $T_{mNN}$ (ms) | $T_{adj}$ (ms) |
| --- | --- | --- | --- | --- | --- | --- | --- |
| Cell $(N^2+1)/2$ | 7x7 2-D tissue | 26.0000 | 7.6000 | 26.0000 | 18.4000 | 18.4000 | 4.8000 |
|  | 15x15 2-D tissue | 48.4000 | 7.6000 | 48.4000 | 40.8000 | 40.8000 | 4.8000 |
|  | 31x31 2-D tissue | 93.2000 | 7.6000 | 93.2000 | 85.6000 | 85.6000 | 4.8000 |
| Cell 1 | 7x7 2-D tissue | 7.6000 | 26.0000 | 42.8000 | 18.4000 | 16.8000 | 4.8000 |
|  | 15x15 2-D tissue | 7.6000 | 48.4000 | 87.6000 | 40.8000 | 39.2000 | 4.8000 |
|  | 31x31 2-D tissue | 7.6000 | 93.2000 | 177.2000 | 85.6000 | 84.0000 | 4.8000 |
| Cell $N^2$ | 7x7 2-D tissue | 42.8000 | 26.0000 | 7.6000 | 16.8000 | 18.4000 | 4.8000 |
|  | 15x15 2-D tissue | 87.6000 | 48.4000 | 7.6000 | 39.2000 | 40.8000 | 4.8000 |
|  | 31x31 2-D tissue | 177.2000 | 93.2000 | 7.6000 | 84.0000 | 85.6000 | 4.8000 |
| Cell 1 to N | 7x7 2-D tissue | 7.6000 | 22.0000 | 36.4000 | 14.4000 | 14.4000 | 4.8000 |
|  | 15x15 2-D tissue | 7.6000 | 41.2000 | 74.8000 | 33.6000 | 33.6000 | 4.8000 |
|  | 31x31 2-D tissue | 7.6000 | 79.6000 | 151.6000 | 72.0000 | 72.0000 | 4.8000 |

Through these tabulations, we found that *T*_*adj*_ is the same despite the alterations in size and the dimension of the tissue when we subject it to D=0.0625 and step time=0.4 ms. Table.9 shows that no *T*_*adj*_ exists when we make the diffusion constant D lesser than 0.0625. Hence, we assign 0.0625 as the minimum diffusion constant, *D*_*min*_. At *D*_*min*_, we already found that *T*_*adj*_ was equal to 4.8 ms. When *D*_*min*_ is doubled, we can infer that *T*_*adj*_ reduces by 2.8 ms. The *T*_*adj*_ at *D*_*min*_ is triple the values of *T*_*adj*_ found when diffusion constant is 3×*D*_*min*_ and 4x*D*_*min*_. After 5×*D*_*min*_, the value of *T*_*adj*_ is constant at 1.2 ms because the slope of the upstroke dominates the dynamics.

**Table 9.** The variation of the relative adjacent peak time, *T*_*adj*_ when the diffusion constant, D is varied according to *D*_*min*_=0.0625. The step time is 0.4 ms.

| Diffusion constant, $D$ | $T_{adj}$ (ms) |
| --- | --- |
| $< D_{min}$ | - |
| $D_{min}$ | 4.8000 |
| $2 \times D_{min}$ | 2.0000 |
| $3 \times D_{min}$ | 1.6000 |
| $4 \times D_{min}$ | 1.6000 |
| $5 \times D_{min}$ | 1.2000 |
| $6 \times D_{min}$ | 1.2000 |
| $7 \times D_{min}$ | 1.2000 |
| $8 \times D_{min}$ | 1.2000 |
| $9 \times D_{min}$ | 1.2000 |
| $10 \times D_{min}$ | 1.2000 |

Fig 17 showcases the propagation wavefront across 4 major junctures, i.e., at cell 1, cell (*N* ^2^ + 1)*/*2, i.e., 113, cell *N* ^2^=225 and from cells 1 to N in the vertical direction where N=15. Fig 17 (A) illustrates the wavefront propagation of a 15×15 2-D FM tissue when cell 1 is stimulated. We can observe that the wavefront propagates diagonally and gradually stimulates the adjacent cells. This propagation occurs till cell *N* ^2^=225. As the APD state of each cell is completed, we can see that the cells within the 2-D FM tissue gradually come to rest. Afterward, we stimulated the cell (*N* ^2^ + 1)*/*2, and the wavefront propagation is displayed in Fig 17 (B). The wavefront is observed to propagate outwardly, stimulating all the cells adjacent to the cell stimulated initially. As the APD phase of all the cells is completed in sequence, the cells in the 2-D tissue gradually come to rest. When the cell *N* ^2^, i.e., 225, is stimulated, we acquire the wavefront propagation shown in Fig 17 (C). We can see that the wavefront propagation is the antithesis of the wavefront propagation found in Fig 17 (A). The wavefront propagates diagonally, stimulating the surrounding cells from cell 225 to cell 1. After completing the APD state, all the tissue cells gradually go to the resting state. As cells 1 to 15 are stimulated in the vertical direction, the wavefront propagation is found to be as illustrated in Fig 17 (D). The wavefront linearly propagates, stimulating all the cells within the 2-D FM tissue. Afterward, we can observe that all the cells gradually move to the resting state. Similar inferences were found for a 7×7 and 31×31 2-D FM tissue.

**Fig 17.**
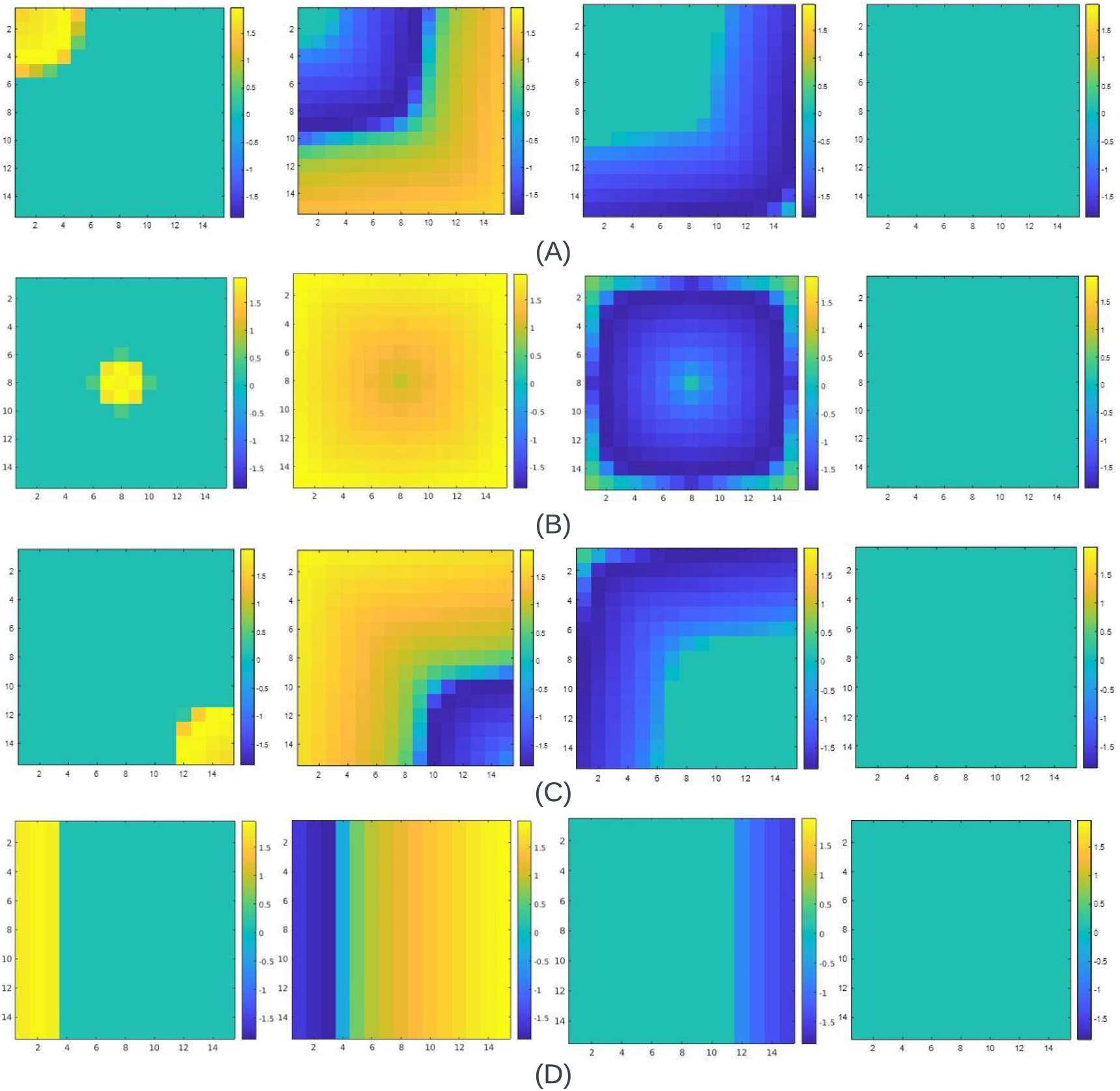
The propagation wavefront of a 15×15 2-D FM tissue of the FHN model when the (A) Cell 1 is stimulated (B) Cell (*N* ^2^+1)/2 is stimulated, i.e., Cell 113 for a 15×15 cell matrix is stimulated (C) Cell NxN, i.e., 225 is stimulated and (D) Multiple cells from Cells 1 to N are stimulated, and N=15 is stimulated. The propagation wave-front starts differently for each stimulation, but in the end, all the cells in the 2-D when the cell or cells are stimulated to go back to rest, tissue goes to rest. The diffusion constant, D used in all cases is 0.00375, and the step time is 0.1 ms.

To create the wavefront propagation of 2-D FM tissue for the FHN model, we utilized the diffusion constant D as 0.00375. We calculated the metrics for all conditions using this value for a step time of 0.1 ms. Table.10 showcases all the parameters above when the cell (*N* ^2^ + 1)*/*2 is stimulated. The *T*_*adj*_ values remain the same through the tabulation. 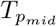 is the same despite the alterations in the size of the 2-D tissue, conveying that the cell (*N* ^2^ + 1)*/*2 is stimulated in all the tissue models. In all 2-D model sizes, we can observe that *T*_*p*1_ and *T*_*pNN*_ are found to be the same. Moreover, *T*_*mc*1_ and *T*_*mNN*_ are also found to be similar, showcasing the outward propagation of the wavefront when the cell (*N* ^2^ + 1)*/*2 is stimulated. Table.10 also showcases all the metrics when cell 1 is stimulated for all the 2-D model sizes. In this case, we can see that *T*_*p*1_ is the same despite the alteration in the 2-D model size. The *T*_*adj*_ is similar in all attempted conditions.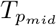, *T*_*pNN*_, *T*_*mc*1_ and *T*_*mNN*_ is found to double when the size of the 2-D tissue alters from 7×7 to 15×15. These almost double when the size alters from 15×15 to 31×31. When cell *N* ^2^ is stimulated, we acquire the metric as tabulated in Table.10. In this case, *T*_*pNN*_ is the same across all the model sizes discussed. This observation shows that the wavefront propagates when cell *N* ^2^ is stimulated. Interestingly, the values of *T*_*p*1_ are equal to the *T*_*pNN*_ and vice-versa when cell 1 is excited. All the values of 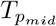 are similar to the ones found when cell 1 is stimulated. *T*_*mc*1_ is found to the same as that of *T*_*mNN*_ as cell 1 is excited. Similar observations can be found when we compare the *T*_*mNN*_ values to *T*_*mc*1_ to the cell 1 stimulation. Overall, this tabulation proves that the wavefront propagation when cell *N* ^2^ is the antithesis of the wavefront propagation when cell 1 is stimulated. Moreover, it also showcases the metrics when a set of cells from 1 to N is stimulated vertically across different 2-D model sizes. We can see that the values of *T*_*adj*_ and *T*_*p*1_ are the same across all the 2-D tissues discussed. We also observed that *T*_*mc*1_ and *T*_*mNN*_ are the same in 7×7,15×15 and 31×31 2-D tissue.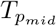 and *T*_*pNN*_ tend to double when the 2-D tissue size alters from 7×7 to 15×15. These tend to double further when the tissue size is enlarged to 31×31.

**Table 10.** The parameters of time for 7×7,15×15 and 31×31 2-D FHN tissue when the diffusion constant, D=0.00375 for stimulations at various junctures. The step time is 0.1 ms.

| Stimulation | 2-D model size | $T_{p_1}$ (ms) | $T_{p_{mid}}$ (ms) | $T_{p_{NN}}$ (ms) | $T_{mc1}$ (ms) | $T_{m_{NN}}$ (ms) | $T_{adj}$ (ms) |
| --- | --- | --- | --- | --- | --- | --- | --- |
| Cell $(N^2 + 1)/2$ | 7x7 2-D tissue | 10.4000 | 2.5000 | 10.4000 | 7.9000 | 7.9000 | 2.0000 |
|  | 15x15 2-D tissue | 19.6000 | 2.5000 | 19.6000 | 17.1000 | 17.1000 | 2.0000 |
|  | 31x31 2-D tissue | 38.0000 | 2.5000 | 38.0000 | 35.5000 | 35.5000 | 2.0000 |
| Cell 1 | 7x7 2-D tissue | 2.5000 | 10.4000 | 17.3000 | 7.9000 | 6.9000 | 2.0000 |
|  | 15x15 2-D tissue | 2.5000 | 19.6000 | 35.7000 | 17.1000 | 16.1000 | 2.0000 |
|  | 31x31 2-D tissue | 2.5000 | 38.0000 | 72.5000 | 35.5000 | 34.5000 | 2.0000 |
| Cell $N^2$ | 7x7 2-D tissue | 17.3000 | 10.4000 | 2.5000 | 6.9000 | 7.9000 | 2.0000 |
|  | 15x15 2-D tissue | 35.7000 | 19.6000 | 2.5000 | 16.1000 | 17.1000 | 2.0000 |
|  | 31x31 2-D tissue | 72.5000 | 38.0000 | 2.5000 | 35.5000 | 34.5000 | 2.0000 |
| Cells 1 to N | 7x7 2-D tissue | 2.5000 | 8.5000 | 14.5000 | 6.0000 | 6.0000 | 2.0000 |
|  | 15x15 2-D tissue | 2.5000 | 16.5000 | 30.5000 | 14.0000 | 14.0000 | 2.0000 |
|  | 31x31 2-D tissue | 2.5000 | 32.5000 | 62.5000 | 30.0000 | 30.0000 | 2.0000 |

Throughout all these analyses, we utilized the diffusion constant, D=0.00375, and we found *T*_*adj*_ to be the same for the various conditions explained in the tabulations. On reducing this value further, we found that the cell-to-cell propagation does not occur when D *<* 0.003673. Hence for Table.11, we took the minimum diffusion constant, *D*_*min*_ as 0.003673. We found that *T*_*adj*_ was 0.5 ms higher than that of the *T*_*adj*_ found when D=0.00375. As the value of D is twice that of *D*_*min*_, we can see that *T*_*adj*_ gets reduced by 1.5 ms. *T*_*adj*_ reduces further by 0.2 ms when D is thrice that of *D*_*min*_. At 4x*D*_*min*_ and 5×*D*_*min*_, the *T*_*adj*_ remains the same, i.e., 0.6 ms. *T*_*adj*_ reduces by 0.1 ms when D=6x*D*_*min*_, D=7×*D*_*min*_ and D=8x*D*_*min*_. At D=9x*D*_*min*_ and D=10x*D*_*min*_, *T*_*adj*_ further reduces by another 0.1 ms. This tabulation infers that *T*_*adj*_ is inversely proportional to D.

**Table 11.** The variation of the relative adjacent peak time, *T*_*adj*_ when the diffusion constant, D is varied according to *D*_*min*_=0.003673. The step time is 0.1 ms.

| Diffusion constant, $D$ | $T_{adj}$ (ms) |
| --- | --- |
| $< D_{min}$ | - |
| $D_{min}$ | 2.5000 |
| $2xD_{min}$ | 1.0000 |
| $3xD_{min}$ | 0.8000 |
| $4xD_{min}$ | 0.6000 |
| $5xD_{min}$ | 0.6000 |
| $6xD_{min}$ | 0.5000 |
| $7xD_{min}$ | 0.5000 |
| $8xD_{min}$ | 0.5000 |
| $9xD_{min}$ | 0.4000 |
| $10xD_{min}$ | 0.4000 |

The evolution of the wavefront propagation is illustrated in Fig 18 for the Fabbri et al. [11] model. When cell 1 is stimulated, we acquire the wavefront propagation as per the illustrations depicted in Fig 18 (A). As cell 1 is stimulated, the adjacent cells get excited, and the wavefront propagates divergently towards cell *N* ^2^, i.e., 225. However, the wavefront also converges towards cell 1. This pattern occurs continuously till the specified period. As a cell, (*N* ^2^ + 1)*/*2=113 is stimulated for a 15×15 2-D tissue; we acquire the wavefront propagation depicted in Fig 18 (B). Cell 113, on stimulation, excites the other adjacent cells, and the wavefront initially propagates outwardly. However, the wavefront is found to propagate in the inwards direction, stimulating the cells that were excited already. This inward propagation occurs continuously till the mentioned time instant. We can observe the opposite wavefront propagation of Fig 18 (A) in the illustrations depicted in Fig 18 (C). As per the illustration, the wavefront propagates outwardly from cell 225 to all the nearby cells. Even before cell 1 is excited, the wavefront is found to be propagating inwardly. This propagation occurs in a recurring order till the last time instant. A linear wavefront propagation occurs continuously when cell 1 to 15 is stimulated. Initially, the wavefront propagates linearly forward. However, as the wavefront propagates forward, the wavefront propagates backward to excite the cells in the 2-D tissue that were excited before. This wavefront propagation is illustrated in Fig 18 (D).

**Fig 18.**
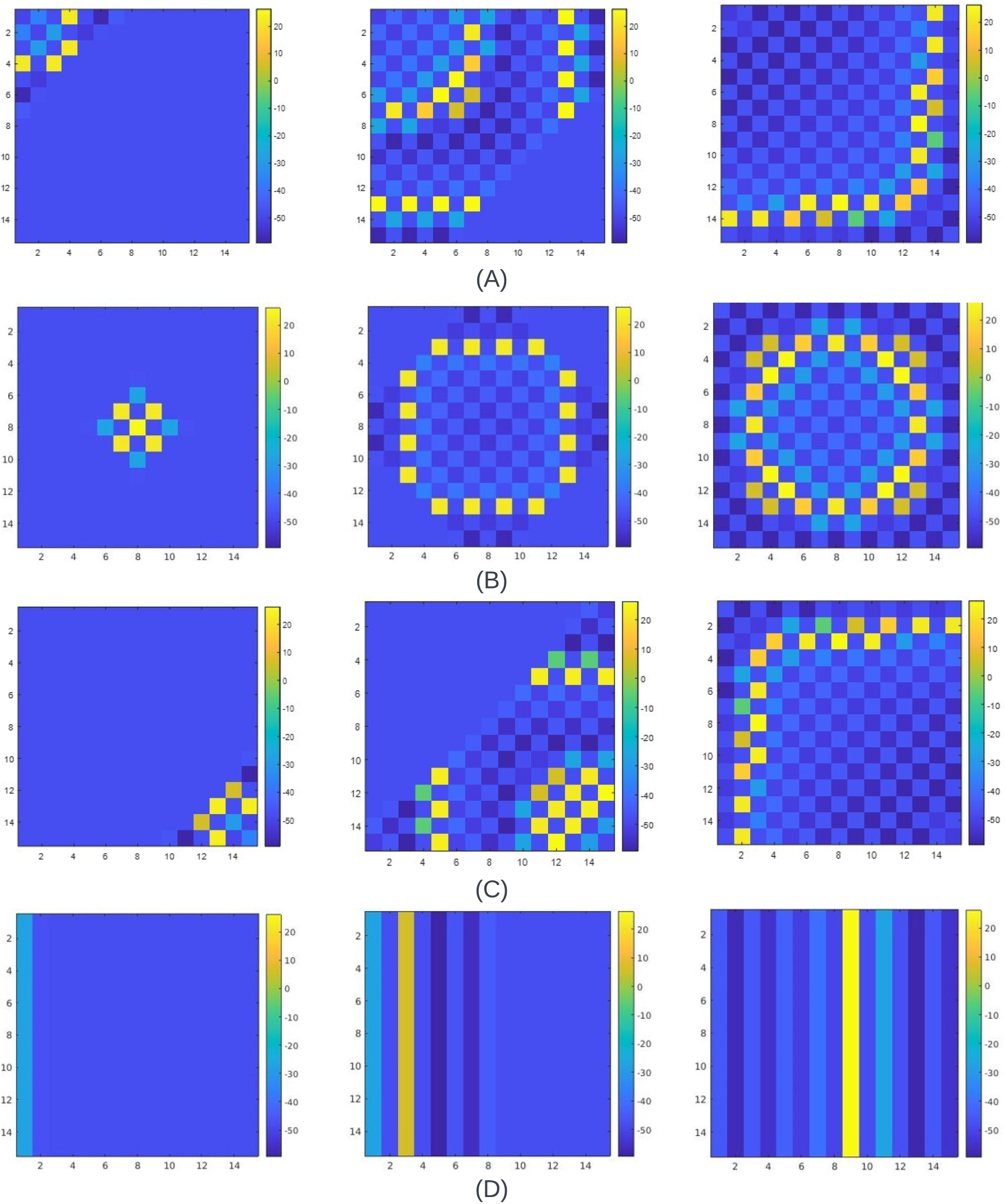
The propagation wavefront of a 15×15 2-D FM tissue of the Fabbri et al. [11] model when the (A) Cell 1 is stimulated (B) Cell (*N* ^2^ + 1)*/*2 is stimulated, i.e., Cell 113 for a 7×7 cell matrix is stimulated (C) Cell NxN, i.e., 225 is stimulated and (D) Multiple cell from Cell 1 to N is stimulated where N=15 is stimulated. The propagation wave-front starts differently for each stimulation and continues to propagate inwardly towards the initial set of cells stimulated. The diffusion constant, D used in all cases is 0.1052, and the step time is 0.02 ms.

Since Fabbri et al. [11] comprises autorhythmic cells, various parameters like the other 2-D models discussed require more investigation. The state controller can be modified to incorporate dynamic properties to replicate behaviors using the S1-S2 protocol [57]. Overall, the illustrations of the wavefront propagations for all the models show that the FM model works as a 2-D tissue.

### Effects of cellular dysfunctions

Previously, we discussed various parameters in terms of time that can be used to justify a 2-D FM tissue. This part will explore how these parameters vary when certain cellular dysfunctions are introduced to the 2-D tissue. A 17×17 2-D FM tissue is created using O’Hara [29], FHN and Fabbri [11] cell models. The 17×17 2-D FM tissues are arranged in a systematic pattern. The top-left corner of the 2-D tissue is cell 1. At the same time, the bottom-right corner of the tissue is a cell *N* ^2^. The middle cell uses (*N* ^2^ + 1)*/*2, i.e., N=145.

Fig 19 illustrates the wavefront propagation for a 17×17 2-D with and without cellular dysfunctions. The stimulus is exerted on the cell 145 to understand the variations in the wavefront propagation with and without cellular dysfunction. Fig 19 (A) shows the wavefront propagation of a 17×17 2-D FM tissue when there is no cellular dysfunction. The wavefront evolves outwardly, as observed in the previous sections. In Fig 19 (B), we induce specific cellular dysfunctions in cells beneath the cell 145. In this case, the wavefront does not propagate through these cells. The wavefront propagates to the other functional cells but bypasses the cells with dysfunctions. As cell 145 goes to rest, all the tissue within the 2-D FM tissue goes to the resting state. We introduce two sets of cellular dysfunctions, and the wavefront propagation is illustrated in Fig 19 (C). In this case, the wavefront bypasses the dysfunctional set of cells above and below the cell 145. The wavefront evolves as usual in the other cells within the 2-D FM tissue. As each cell within the 2-D FM tissue goes to the resting state, the entire set of cells, barring those with dysfunctions, are found to go towards the resting state gradually.

**Fig 19.**
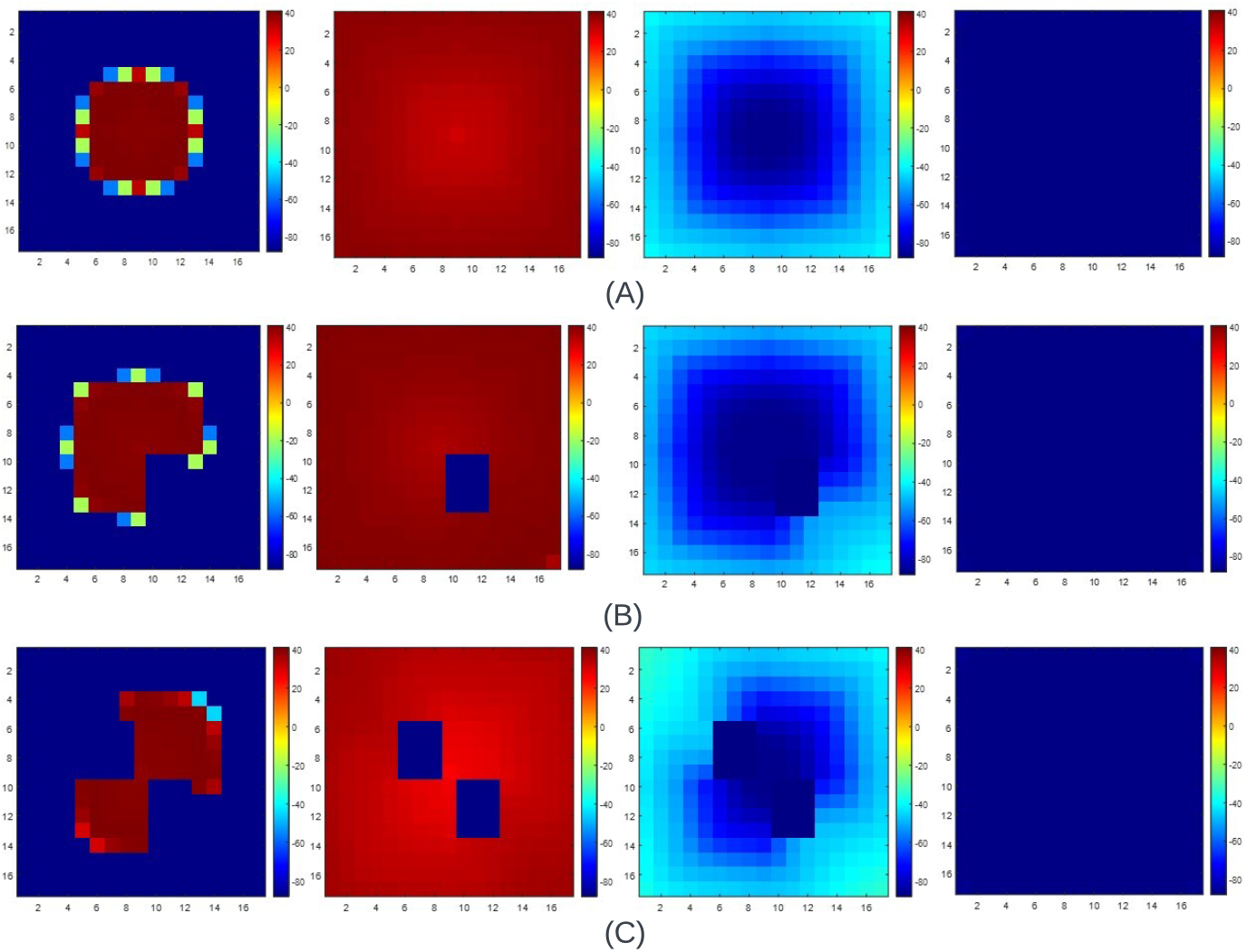
The propagation wavefront of a 17×17 2-D FM tissue when the cell, (*N* ^2^ + 1)*/*2=145, is stimulated when there are (A) no dysfunctions, (B) One dysfunction, (C) Two dysfunction. The diffusion constant, D, used in all cases is 0.0625, and the step time is 0.4 ms.

The tabulations of these illustrations are depicted in Table.12. The wavefront without the cellular dysfunction is observed to have the same value for *T*_*p*1_ and *T*_*pNN*_ which in turn leads to the values of *T*_*mc*1_ and *T*_*mNN*_ to be equal. However, when the dysfunctions in these cells are introduced, as per Fig 19 (B), we can observe that there is a surge in the values of *T*_*pNN*_ and *T*_*mNN*_ by 6.4 ms. When two sets of cellular dysfunctions exist, as in Fig 19 (B), we can see the overall increase in the values of *T*_*p*1_ and *T*_*pNN*_, which in turn leads to the surge in the values of *T*_*mc*1_ and *T*_*mNN*_. However, the values of 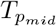 and *T*_*adj*_ remain the same in all the cases. Overall, there is some drastic alteration in the mentioned parameters when certain cellular dysfunctions are introduced. We then look to comprehend the wavefront propagation using the FHN model in a 2-D FM tissue where similar cellular dysfunction can be done.

**Table 12.** The time parameters for 17×17 2-D FM tissue using different cell models with diffusion constant, D and step time when the middle cell (*N* ^2^+1)/2=145 is stimulated with and without cellular dysfunctions.

| Cell model | D | Step Time (ms) | Dysfunction | $T_{p1}$ (ms) | $T_{pmid}$ (ms) | $T_{pNN}$ (ms) | $T_{mcl}$ (ms) | $T_{mNN}$ (ms) | $T_{adj}$ (ms) |
| --- | --- | --- | --- | --- | --- | --- | --- | --- | --- |
| O'Hara [29] | 0.0625 | 0.4 | None | 54.0000 | 7.6000 | 54.0000 | 46.4000 | 46.4000 | 4.8000 |
|  |  |  | One set | 54.0000 | 7.6000 | 60.4000 | 46.4000 | 52.8000 | 4.8000 |
|  |  |  | Two sets | 62.0000 | 7.6000 | 60.4000 | 54.4000 | 52.8000 | 4.8000 |
| FHN | 0.003673 | 0.1 | None | 25.6000 | 2.5000 | 25.6000 | 23.1000 | 23.1000 | 2.5000 |
|  |  |  | One set | 25.6000 | 2.5000 | 29.4000 | 23.1000 | 26.9000 | 2.5000 |
|  |  |  | Two sets | 30.3000 | 2.5000 | 29.4000 | 27.8000 | 26.9000 | 2.5000 |

Fig 20 depicts the wavefront propagation of the FHN model using a 17×17 2-D FM tissue. As mentioned earlier, the minimum diffusion constant, *D*_*min*_, was found to be 0.003673. Hence, we will utilize this value for a time step of 0.1 ms. The wavefront evolves outwardly in all directions when no cellular dysfunction exists, as depicted in Fig 20 (A). When cellular dysfunction exists, as in Fig 20 (B), we can observe that the wavefront propagation tends to elude these cells. The other functional cells tend to propagate outwardly, and all of them are found to move gradually toward the resting state when there are two cellular dysfunctions, as shown in Fig 20 (C), the wavefront propagates by eluding these areas. As another cell, 145, goes to rest, all the functional cells surrounding it go to the resting state. From this illustration, we can comprehend that the wavefront propagation evolves differently whenever a set of cells is dysfunctional.

**Fig 20.**
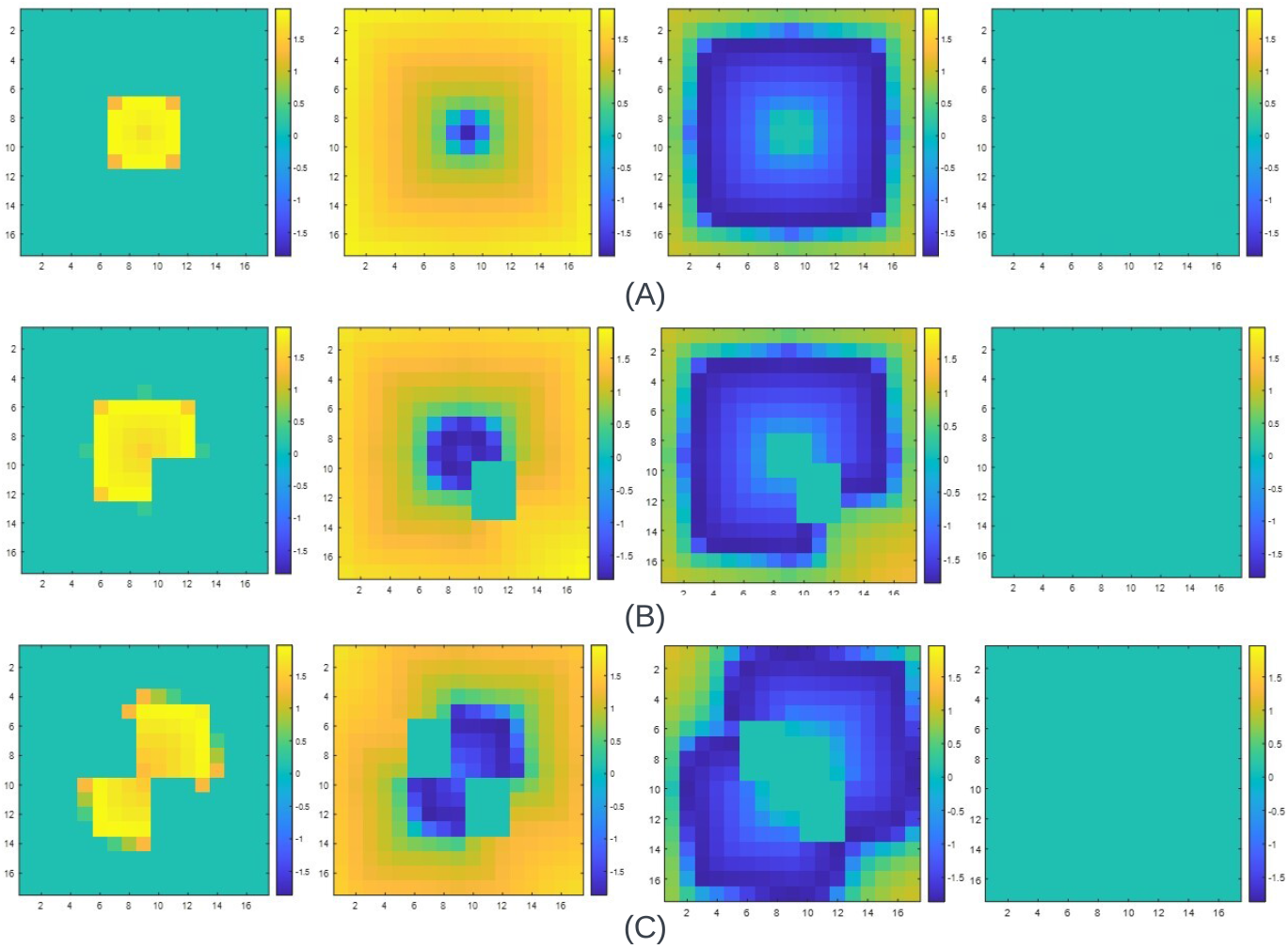
The propagation wavefront of a 17×17 2-D FM tissue using FHN model when the cell (*N* ^2^ + 1)*/*2=145 is stimulated when there are (A) no dysfunctions (B) One dysfunction (C) Two dysfunction. The diffusion constant, D, used in all cases is 0.003673, and the step time is 0.1 ms.

Without introducing certain dysfunctions to the 2-D FM model using the FHN AP characteristics, we can observe that the parameters *T*_*adj*_ and 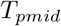 remain the same (refer Table.12). The values of *T*_*p*1_ and *T*_*pNN*_ are the same, which in turn leads to the parameters *T*_*mc*1_ and *T*_*mNN*_ being similar. We can observe that the parameter *T*_*pNN*_ tends to surge by 3.8 ms, leading to the alteration in the value of *T*_*mNN*_ by the same margin as dysfunctions are introduced as in Fig 20 (B). In this case, the value of *T*_*p*1_ and 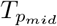 is the same as the wavefront propagation in Fig 20 (A). As cellular dysfunctions are introduced in Fig 20 (C), we can observe that the values of *T*_*p*1_ and *T*_*pNN*_ tend to rise, which leads to the rise in the values of *T*_*mc*1_ and *T*_*mNN*_. We can see that certain parameters tend to increase when cellular dysfunctions occur in the 2-D FM tissues using the FHN cell model.

The illustrations in Fig 21 show the wavefront propagation using the autorhythmic cells for different conditions. In all the cases, we have stimulated cell 145. Fig 21 (A) illustrates the wavefront propagation of 17×17 2-D tissues using the pacemaker cell when no cellular dysfunctions exist. We can observe that the wavefront initially propagates outwardly and later evolves inwardly towards the cell 145. It propagates inwardly towards cell 145 until the last instant. When a set of cells below the cell 145 becomes dysfunctional, the propagation wavefront is depicted in Fig 21 (B). We can see that throughout the propagation wavefront, all the other cells, barring the cells with dysfunctions, get stimulated, and the inward pattern towards the cell 145 prolongs the required time frame. When two sets of cells are dysfunctional above and below cell 145, we get the wavefront propagation as in Fig 21 (C). In this case, the wavefront bypasses the dysfunctional set of cells to propagate inwardly towards the cell 145. These illustrations showed the propagation wavefront in the 2-D FM tissue with and without dysfunction. Further investigation into the S1-S2 protocol and APD restitution [57] is required to calculate all the time parameters like the other 2-D tissue models. Overall, cellular dysfunctions affect the wavefront propagation in a 2-D for different cells.

**Fig 21.**
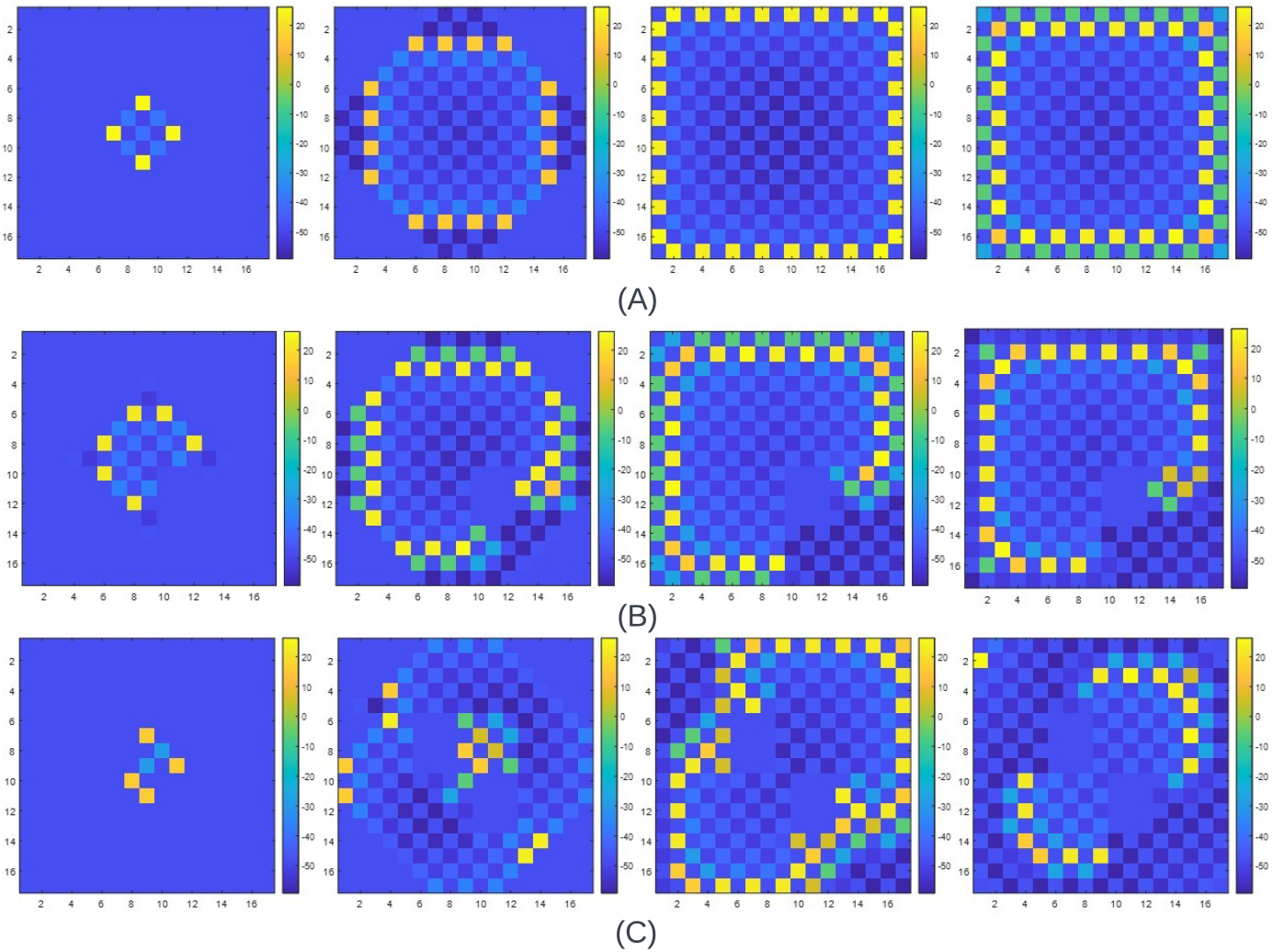
The propagation wavefront of a 17×17 2-D FM tissue using Fabbri et al. [11] model when the cell (*N* ^2^ + 1)*/*2=145 is stimulated when there are (A) no dysfunctions (B) One dysfunction (C) Two dysfunction. The diffusion constant, D, used in all cases is 0.1052, and the step time is 0.02 ms.

## Discussion

This study presents a novel FM model designed to alter the characteristics of a single sine wave based on the APDs of a cell model: Beeler Reuter (BR), Fenton Karma (FK), FitzHugh Nagumo and Fabbri et al. [11] model. Before evaluating how precisely this novel methodology can replicate the APD, various test signals were used to comprehend the efficacy of this method. Table.1 and Table.2 tabulates various values, which showcases how this model effectively replicates the test signals.

Afterward, we reconstruct the Δ_*w*_ and APD waveshapes for the four cell models to further confirm the model’s effectiveness. The RMSE metrics in Table.3 are absolute measurements. The percentage error for the Δ_*w*_ and the APD for the BR model are 1.28% and 3.6%, respectively. The closeness of the traces in Fig. 4 and 5 reflect these percentage errors. The high *R*^2^ metrics for Δ_*w*_ and the APD reconstructions confirm the fidelity of the reconstructed signals using the FM methodology. Notably, the FM model can capture any dynamic changes in the morphologies by adapting the frequency modulating factor, Δ_*w*_. Hence, this model can potentially simulate the effects of new drugs with the *I*_*f*_ blockade, 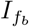 (refer Table.4).

Utilizing a piecewise linear segment reduces the Look-Up-Table (LUT) size from K samples to n segments. For example, the Δ_*w*_ profile has 40000 samples for the BR model and would require an LUT with 40000 elements. Using piecewise linear fitting, the LUT size equals the number of segments (refer Table.3).

We introduced a state controller to the FM model to facilitate the interconnection of cells and, hence, implement various tissue models. Figures showing wavefront propagation for 1-D and 2-D tissues have been presented.

This study shows the successful simulation of the proposed modeling approach to replicate a cell model’s AP precisely. Although dynamic properties, like APD restitution and action potential rate dependence, have not been included in this study, the state controller can be enhanced (as has been done for 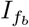) to incorporate these properties. Given the generic nature of this model, it can be expanded to various biological cells like skeletal muscle cells, atrial myocytes, plant cells, and even the cells of diverse species.

The proposed novel FM model has the potency for a low-cost computational approach that may allow researchers to explore new avenues. This area requires an in-depth investigation to guarantee maximum effectiveness and efficiency. Moreover, the fitting piecewise polynomial can be altered from linear to quadratic or cubic to reduce the number of breakpoints, which may provide a more accurate APD waveshape.

## Conclusion

The proposed FM model needs to be reported in the literature. It accurately captures the electrical activities of various types of biological cells. It replicates the action potential durations (APDs) and precisely accounts for the effects in the blockage current. The performance of this model closely aligns with biophysical simulations and clinical data.

The model is inherently simple and holds significant potential for emulating multiple cells, thereby performing tissue-level simulations. Implementing this model is straightforward and enables a seamless application to various fields where processes exhibit (non-linear) periodic or quasi-periodic characteristics. This adaptability can facilitate advancement in various research domains and enhance real-time emulations, especially on an FPGA.

## Supporting Information

**S1 Fig. The nine sine waves utilized for Test Case 2**. The illustration of nine sine waves with varying amplitude and frequency in Table.2.

**S2 Fig. Minimum piecewise linear fits for the polynomial values of** Δ_*w*_ **of the sinoatrial node cell at different levels of** *I*_*f*_ **blockade**. The red dots represent the each part of Δ_*w*_ when the *I*_*f*_ blockade,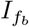 is varied from 0 to 1. The dotted blue lines show the trajectory of the fitted linear polynomial. *ω* is the fundamental frequency and the parts of Δ_*w*_ are represented as coefficients A1,A2,A3,A4,., A29.

**S1 Movie. Propagation wavefront of a 17×17 2-D tissue of the O’Hara [29] model when the stimulation is done on cell, N=**(*N* ^2^ + 1)*/*2 **where diffusion constant, D=0.0625 and step time is 0.4 ms**. Movie illustrates the propagation wavefront of a 17×17 2-D tissue of the O’Hara [29] model when cell N=(*N* ^2^ + 1)*/*2 is stimulated.

**S2 Movie. Propagation wavefront of a 17×17 2-D tissue of the FHN model when the stimulation is done on cell, N=**(*N* ^2^ + 1)*/*2. **Diffusion constant, D is 0.003673 and step time is 0.1 ms**. Movie illustrates the propagation wavefront of a 15×15 2-D tissue of the FHN model when cell N=(*N* ^2^ + 1)*/*2 is stimulated.

**S3 Movie. Propagation wavefront of a 17×17 2-D tissue of the Fabbri [11] model when the stimulation is done on cell, N=**(*N* ^2^ + 1)*/*2. **Diffusion constant, D is 0.1052 and step time is 0.02 ms**. Movie illustrates the propagation wavefront of a 15×15 2-D tissue of the Fabbri [11] model when cell N=(*N* ^2^ + 1)*/*2 is stimulated.

**S4 Movie. The propagation wavefront of a 17×17 2-D FM tissue using O’Hara [29] model when the cell, N=**(*N* ^2^ + 1)*/*2**=145 is stimulated when there is one set of cellular dysfunction. Diffusion constant, D is 0.0625 and step time is 0.4 ms**. Movie shows how the propagation wavefront for a 17×17 2-D tissue alters when there is a set of cellular dysfunctions.

**S5 Movie. The propagation wavefront of a 17×17 2-D FM tissue using O’Hara [29] when the cell, N=**(*N* ^2^ + 1)*/*2**=145 is stimulated when there are two sets of cellular dysfunction. Diffusion constant, D is 0.0625 and step time is 0.4 ms**. Movie shows how the propagation wavefront for a 17×17 2-D tissue alters when there are two sets of cellular dysfunctions.

**S6 Movie. The propagation wavefront of a 17×17 2-D FM tissue using FHN model when the cell, N=**(*N* ^2^ + 1)*/*2**=145 is stimulated when there is one set of cellular dysfunction. Diffusion constant, D is 0.003673 and step time is 0.1 ms**. Movie shows how the propagation wavefront for a 17×17 2-D tissue alters when there is a set of cellular dysfunctions.

**S7 Movie. The propagation wavefront of a 17×17 2-D FM tissue using FHN model when the cell, N=**(*N* ^2^ + 1)*/*2**=145 is stimulated when there are two sets of cellular dysfunction. Diffusion constant, D is 0.003673 and step time is 0.1 ms. Diffusion constant, D is 0.003673 and step time is 0.1 ms**. Movie shows how the propagation wavefront for a 17×17 2-D tissue alters when there are two sets of cellular dysfunctions.

**S8 Movie. The propagation wavefront of a 17×17 2-D FM tissue using Fabbri [11] model when the cell, N=**(*N* ^2^ + 1)*/*2**=145 is stimulated when there is one set of cellular dysfunction. Diffusion constant, D is 0.1052 and step time is 0.02 ms**. Movie shows how the propagation wavefront for a 17×17 2-D tissue alters when there is a set of cellular dysfunctions.

**S9 Movie. The propagation wavefront of a 17×17 2-D FM tissue using Fabbri [11] model when the cell, N=**(*N* ^2^ + 1)*/*2**=145 is stimulated when there are two set of cellular dysfunction. Diffusion constant, D is 0.1052 and step time is 0.02 ms**. Movie shows how the propagation wavefront for a 17×17 2-D tissue alters when there are two sets of cellular dysfunctions.

